# Phylogenomics of piranhas and pacus (Serrasalmidae) uncovers how convergent diets obfuscate traditional morphological taxonomy

**DOI:** 10.1101/2020.03.02.973503

**Authors:** M.A. Kolmann, L.C. Hughes, L.P. Hernandez, D. Arcila, R. Betancur, M.H. Sabaj, H. López-Fernández, G. Ortí

## Abstract

The Amazon and neighboring South American river basins harbor the world’s most diverse assemblages of freshwater fishes. One of the most prominent South American fish families are the Serrasalmidae (pacus and piranhas), found in nearly every continental basin. Serrasalmids are keystone ecological taxa, being some of the top riverine predators as well as the primary seed dispersers in the flooded forest. Despite their widespread occurrence and notable ecologies, serrasalmid evolutionary history and systematics are controversial. For example, the sister taxon to serrasalmids is contentious, the relationships of major clades within the family are obfuscated by different methodologies, and half of the extant serrasalmid genera are suggested to be non-monophyletic. We used exon capture to explore the evolutionary relationships among 64 (of 99) species across all 16 serrasalmid genera and their nearest outgroups, including multiple individuals per species in order to account for cryptic lineages. To reconstruct the timeline of serrasalmid diversification, we time-calibrated this phylogeny using two different fossil-calibration schemes to account for uncertainty in taxonomy with respect to fossil teeth. Finally, we analyzed diet evolution across the family and comment on associated changes in dentition, highlighting the ecomorphological diversity within serrasalmids. We document widespread non-monophyly within Myleinae, as well as between *Serrasalmus* and *Pristobrycon*, and propose that reliance on traits like teeth to distinguish among genera is confounded by ecological convergence, especially among herbivorous and omnivorous taxa. We clarify the relationships among all serrasalmid genera, propose new subfamily affiliations, and support hemiodontids as the sister taxon to Serrasalmidae.

## INTRODUCTION

The family Serrasalmidae, piranhas and pacus (Fig. 1), is a diverse freshwater clade of characiform fishes found throughout tropical and subtropical South America. Ninety-seven extant species are primarily distributed east of the Andes, with just a single species found west in the Maracaibo Basin, but the fossil record extends the historical distribution of the family further west into the Magdalena region (Lundberg et al., 2010). While piranhas are generally carnivorous, their sister taxa, the pacus, are herbivores that include some of the primary seed dispersers in flooded forests or varzea (Goulding, 1980; Correa et al., 2007). Despite their keystone status and commercial significance throughout Amazonia, the ecologies of piranhas and pacus are poorly understood and often misrepresented. The ferocious reputation of piranhas stems largely from accounts of their feeding on corpses (Sazima & Guimarães, 1987) or nipping bathers, the latter attributed to the protective nature of piranhas defending nests (Haddad & Sazima, 2003, 2010; but see Kolmann et al., 2018b). We have only recently understood the role that pacus play in structuring forests, as large frugivores that can disperse seeds over great distances (ichthyochory) (Correa et al., 2015, 2016). Some rainforest trees appear particularly specialized for ichthyochory, with seeds that have greater germination probability after digestion by fishes (Anderson et al., 2009) or by bearing fleshy fruit with increased buoyancy (Horn et al., 2011; Correa et al., 2018). Large pacus like the iconic tambaqui (*Colossoma*) and pirapitinga (*Piaractus*) are attracted to the sound of fruit falling in the water, even ‘staking out’ fruiting trees for weeks at a time (Goulding, 1980).

**Figure 1.**
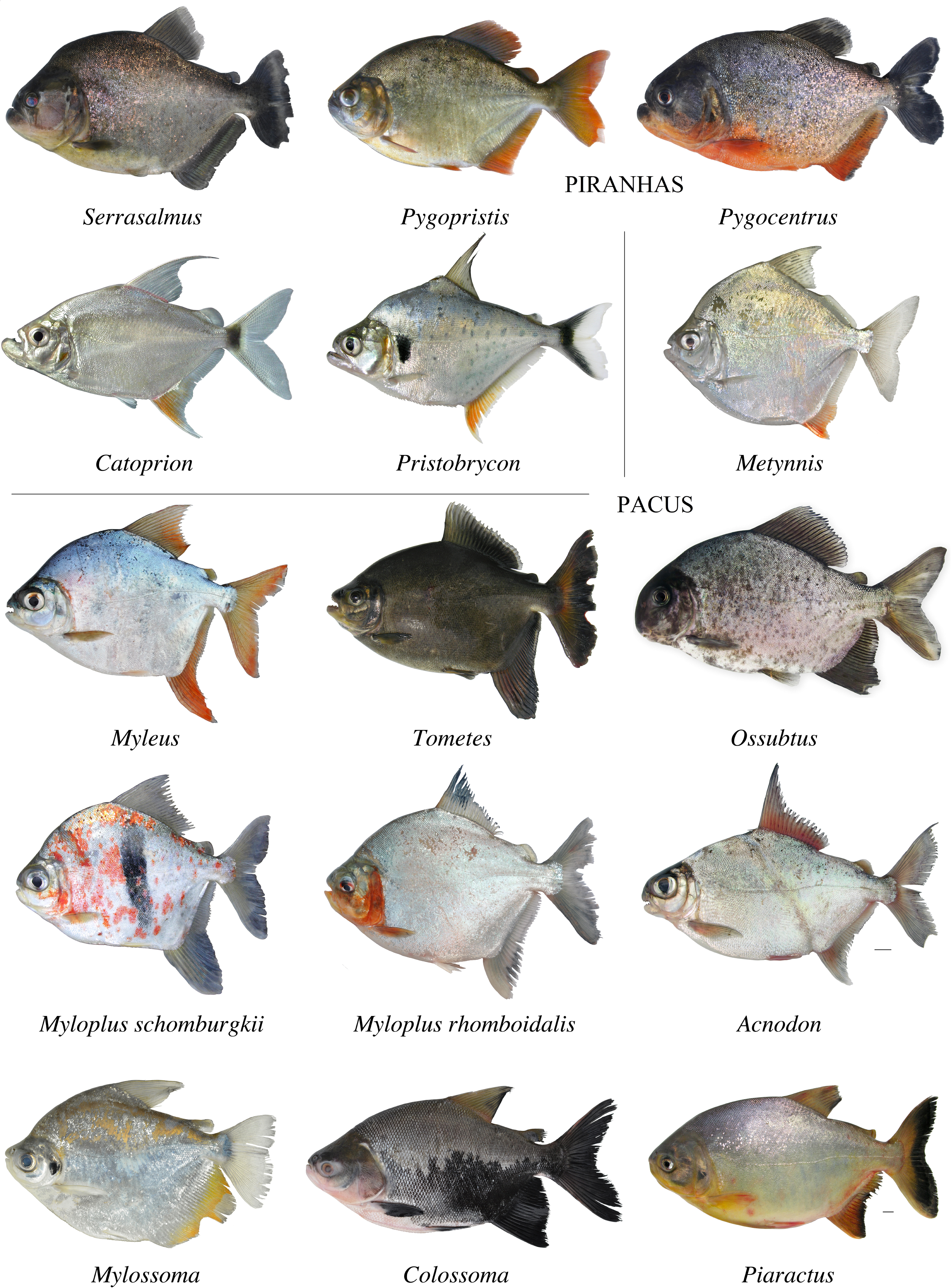
Morphological diversity of various serrasalmid species and genera. Piranhas divided from pacus by vertical and horizontal lines. Photo of *Ossubtus* by L. Sousa, others by M. Sabaj.

From fruits and seeds, to fins and flesh, serrasalmids feed on a wide variety of prey and prey materials (Correa et al., 2007), but many genera remain understudied regarding their evolutionary ecology. Medium-sized pacus (Myleinae) feed heavily on the leaves, flowers, and stems of riparian and aquatic plants (Correa & Winemiller, 2014), with some adapted to scraping river weed (Podostemaceae) off rocks in a manner unique among otophysan fishes (Andrade et al., 2019a; Huie et al., 2019). Contrary to the popular idea of piranhas as exclusively carnivorous, their diets vary greatly across seasons, ontogeny, and species. Some piranhas are facultative frugivores (*Pristobrycon*; Nico & Taphorn, 1988; Correa & Winemiller, 2014), while others feed on either fish scales (lepidophages, i.e. *Catoprion*; Goulding, 1980; Sazima & Machado, 1990; Kolmann et al., 2018a) or fins (pterygophages, i.e. *Serrasalmus elongatus*; Röpke et al., 2014). How such diverse diets have evolved in serrasalmids remains uncertain, despite efforts by authors like Correa et al. (2007), because of the lack of a well resolved phylogenetic framework for the family.

Early classifications (Eigenmann, 1915; Norman, 1929; Gosline, 1951; Géry, 1977) relied on dentition to divide serrasalmids into two major groups, those with two rows of teeth on the upper jaw (‘herbivorous’ pacus) vs. one row (‘carnivorous’ piranhas). A more comprehensive and cladistic analysis of morphology by Machado-Allison (1982, 1983, 1985) also divided serrasalmids into two major groups, but disagreed on their composition by transferring *Metynnis*, previously classified as a pacu, to the piranha clade. The cladistic analysis of the newly discovered fossil †*Megapiranha* by Cione et al. (2009) supported the placement of *Metynnis* sister to piranhas and provided the first morphological evidence for the non-monophyly of the remaining pacus. Cione et al. (2009) and more recent studies have expanded the breadth of ecological and morphological diversity among serrasalmids. For example, some species of pacus have heterodont dentitions which vary along numerous phenotypic axes, from molariform to incisiform, spatulate to crenulate (Huie et al., 2019; Kolmann et al., 2019). In fact, tooth characteristics and arrangement remain the best characters for distinguishing pacu taxa (Andrade et al., 2013, 2016; Nico et al., 2018).

Molecular phylogenies (Ortí et al., 1996, 2008) support the morphological phylogeny of Cione et al. (2009) wherein the piranhas + *Metynnis* clade are sister to medium-sized taxa like *Acnodon* and *Myleus* and are therefore nested within pacus (Supplements). However, molecular studies using mitochondrial genes (Ortí et al., 1996, 2008; Hubert et al., 2007; Freeman et al., 2007) revealed problems with generic monophyly and failed to reconcile genus-level relationships. Thompson et al. (2014) constructed the most rigorous serrasalmid phylogeny to date and found rampant non-monophyly within Myleinae and among *Serrasalmus* and *Pristobrycon* species (Supplements). These molecular analyses had limited taxonomic sampling, including only two of the five *Pristobrycon* species, and also lacked *Utiaritichthys*, which DNA barcoding suggests is nested within *Myloplus* (Machado et al. 2018).

Improving our understanding of the timeline of lineage, phenotypic, and ecological diversification in serrasalmids thus requires broader species-level sampling. Resolution of recalcitrant relationships or questionable monophyly (e.g. *Serrasalmus*) should benefit from phylogenomic approaches that leverage much larger datasets than previously available. Thompson et al. (2014) dated their serrasalmid phylogeny with two fossil calibrations and found the basal divergence between pacus and piranhas (+ *Metynnis*) starting during the middle Paleocene (∼ 60 mya) and the diversification of Myleinae starting in late Eocene (45 mya). However, the age of other lineages within serrasalmids are more uncertain, despite a rich fossil record that could improve our understanding of the timeline of diversification within the family and their nearest characiform relatives (Dahdul, 2004, 2007).

We used exon-capture phylogenomic methods with greater taxon sampling than any previous study to reassess relationships among serrasalmid genera. Using multiple fossil calibrations to estimate relative divergence times among lineages, we explored how uncertainty about fossil choice shapes our estimates of the timing of lineage diversification in serrasalmids. Finally, we used ancestral character state estimation to explore how diet diversity, novelty, and convergence evolved across extant serrasalmid lineages, and how major diet specializations have evolved in this clade. We document how ecological convergence in diet may be shaping dental morphology and the effect this phenomenon has on taxonomic character states used to distinguish among serrasalmid genera.

## METHODS

### Taxonomic Sampling

Exon sequence data were analyzed for 194 individuals (Table S1) including 44 previously published in Arcila et al. (2017) and Betancur et al. (2019). Outgroups included 59 individuals representing 50 species distributed among 38 genera in 12 families of Characiformes. The ingroup, Serrasalmidae, included 135 individuals representing all 16 nominal valid genera and at least 65 of the 99 nominal valid species. Twelve specimens were excluded due to potential mislabeling or contamination problems. A probe set for exon capture previously optimized for Otophysi by Arcila et al. (2017) was used to capture 1051 exons for sequencing. Library preparation was performed at Arbor Biosciences (www.arborbiosci.com), using a dual-round hybridization protocol (Li et al. 2013) to enrich the libraries for the targeted exons on pools of eight species. Paired-end sequencing (100 bp) was performed at the University of Chicago Genomics Facility on a HiSeq 4000. Up to 192 enriched libraries were combined to form multiplex pools for sequencing in a single lane.

### DNA extraction and Exon Capture Protocols

Genomic DNA was extracted from muscle biopsies or fin clips preserved in 90-99% ethanol using either Qiagen DNEasy kits or the phenol-chloroform protocol in the Autogen platform available at the Laboratory of Analytical Biology at the National Museum of Natural History (Smithsonian Institution) in Washington, D.C. Laboratory protocols for library preparation and probe sets for exon capture were optimized for Otophysi, and followed the procedures described in Arcila et al. (2017). Library preparation and target enrichment was completed at Arbor Biosciences (www.arborbiosci.com). Libraries were sequenced on an Illumina HiSeq 4000 at the University of Chicago Genomics Facility.

### Data Assembly and Alignment

We used the bioinformatics pipeline optimized by Hughes et al. (2020) to obtain sequence alignments for 951 exon markers from an initial set of 1051 (S1 appendix). Raw FASTQ files were trimmed with Trimmomatic v0.36 (Bolger et al. 2014), to remove low quality sequences and adapter contamination. Trimmed reads were then mapped with BWA-MEM (Li & Durbin, 2009) against a fasta file containing all sequences used for bait design (see Arcila et al 2017). SAMtools v1.8 was used to remove PCR duplicates and sort the reads that mapped to each of the exons (Li et al. 2009b). Sorted reads (by species) were then assembled individually for each exon using Velvet (Zerbino & Birney, 2008), and the longest contig produced by Velvet was used as the initial reference sequence for input to aTRAM v2.0 (Allen et al. 2017). aTRAM was run with Trinity v2.8.5 as the assembler to extend contigs iteratively. Redundant contigs with 100% identity produced by aTRAM were removed with CD-HIT v4.8.1 using CD-HIT-EST (Li & Godzik, 2006; Fu et al., 2012). Open reading frames for remaining contigs were identified with Exonerate (Slater & Birney, 2005). Sequences for each exon were aligned with MACSE v2.03 (Ranwez et al. 2018).

Resulting data matrices were combined with previously published exon sequences (Arcila et al., 2017; Betancur-R. *et al*. 2019) (Table S1). For quality control and assessment, estimated gene trees were visually assessed and flagged when Serrasalmidae was not monophyletic. These gene trees were visually inspected to detect and remove putatively paralogous sequences, and samples with alarmingly long branches. Alignments with fewer than 100 sequences also were removed from downstream analyses.

### Phylogenetic inference

A species tree was estimated under the multi-species coalescent (MSC) using ASTRAL-III v5.6.3 (Mirarab et al., 2014; Zhang et al., 2018), with individual gene trees estimated under maximum likelihood (ML) using IQTREE v1.6.10 (Nguyen et al., 2014). Each locus was partitioned by codon position, according to automatic model selection parameters obtained from ModelFinder using the ‘TESTMERGE’ option (Kalyaanamoorthy et al., 2017), with ten independent Maximum Likelihood searches for each gene alignment. Concatenated amino acid and nucleotide matrices also were analyzed with IQTREE, with nucleotide sequences partitioned by codon position and the best substitution model was fitted using the ‘TEST’ option of ModelFinder. Protein sequences were translated from nucleotides using AliView v1.0 (Larsson, 2014) and the best model across all genes selected using ModelFinder. Ten independent searches were run for each concatenated analysis. Branch support for the ML analyses of concatenated matrices was assessed with 1,000 ultra-fast bootstrap (UFBoot) replicates (Minh et al., 2013) and 1,000 SH-like approximate likelihood ratio test (SH-aLRT) replicates (Guindon et al., 2010). Support for the species tree topology obtained with ASTRAL-III was assessed with local posterior probabilities (PP, Sayyari & Mirarab, 2016).

### Fossil calibration

The oldest fossils associated with Serrasalmidae are isolated pacu-like teeth from the Bolivian El Molino Formation (∼73-60 mya; Gayet, 1991; Gayet & Meunier, 1998). Although often used to date the origin of the family, these fossil teeth are unusual for serrasalmids for several reasons: (1) their small size of < 0.75-1.0 mm (Gayet, 1991; Gayet & Meunier, 1998; Gayet et al., 2001) despite most serrasalmid teeth being far larger (>1 cm; Shellis & Berkovitz, 1976; Kolmann et al., 2019); (2) pacu teeth, while complex in shape, lack the lingual cusp visible in Gayet et al. (2001, Fig. 7d); (3) these fossil teeth lack any of the interlocking morphologies typical of extant serrasalmid dentitions (Kolmann et al., 2019); (4) the timeline of Bolivian serrasalmid fossil teeth leave a 25-32 mya gap in the fossil record until the first confidently-identified serrasalmid tooth (∼38 mya; DeCelles & Morton, 2002; Dahdul, 2007; Lundberg et al., 2010).

The El Molino fossils may represent juvenile serrasalmid teeth, but this hypothesis seems unlikely due to the absence of larger adult teeth and the well-known taphonomic bias towards larger-sized skeletal elements. Furthermore, these fossil teeth strongly resemble dentitions from distantly related alestids (Alestidae) discovered in various North African deposits (Murray et al., 2003, 2004a, 2004b). It is possible that these Bolivian fossils are stem characoids, or simply indistinguishable at a more circumscribed exclusive taxonomic level (as others have suggested, see Patterson, 1993; Otero et al., 2008). This scenario is supported by the non-monophyly of South American characoids with respect to the African Alestoidea (Alestidae+Hepsetidae), and its sister group relation to exclusively Neotropical taxa in Neotropical Erythrinoidea + Curimatoidea (Arcila et al., 2017; Betancur et al., 2019). Given this uncertainty, we excluded the putative pacu teeth from the El Molino formation in our first set of calibrations (Scheme 1) but included them in our second set to assess its effect on divergence times within the family (Scheme 2; Supplemental Materials). These calibration schemes included 11 outgroup fossils, and three additional serrasalmid calibration points are based on Miocene fossils summarized in Lundberg et al. (2010).

As running time-calibration analysis using a Bayesian framework for large phylogenomic datasets can be computationally intensive, we randomly selected four 50-gene subsets of the 200 most complete genes and pruned each subset to include one tip per taxon. We transformed the concatenated ML DNA topology into a chronogram under penalized likelihood using the *chronos* function in R (ape v5.3; Paradis et al., 2019). This chronogram was used as a starting tree for BEAST 2 v2.5.0 (Bouckaert et al., 2014) to generate relaxed-clock divergence time estimates, the topology of the resulting trees was also constrained to match the concatenated nucleotide phylogeny. Each subset of 50 genes was run independently twice for 2.0×10^8^ generations. Convergence was assessed in Tracer v1.7.1 (Rambaut et al., 2018) by checking that ESS values were greater than 200 for all parameters. Independent runs from each of the four different subsets were combined in LogCombiner if their 95% highest posterior densities for divergence times overlapped, and a maximum clade credibility tree was generated in TreeAnnotator for each of the two calibration schemes.

### Ancestral state reconstruction of diet evolution in piranhas and pacus

Serrasalmid taxa were assigned to 11 categories based on diet (Supplemental Materials): (1) omnivores (plant and fish feeders), (2) piscivores, (3) planktivores, (4) plankton & plant materials, (5) plant materials & invertebrates, (6) plant materials, invertebrates & scales, (7) plant materials & seeds (granivores), (8) plant materials, seeds & fruit (frugivores), (9) plant materials, seeds & invertebrates, (10) fish & fish parts, and (11) phytophagous. Fish & fish parts applies to taxa that consume some combination of scales, fin rays, and whole fishes (e.g., *Serrasalmus elongatus*, a fin-nipper or *Catoprion*, a scale-feeder; Gonzalez & Vispo, 2003; Röpke et al., 2014; Nico & Morales, 1994). We used stochastic character mapping (Huelsenbeck et al., 2003; Bollback, 2006) to reconstruct the evolution of these diet modes across the dated phylogeny (Scheme 1) with the *make*.*simmap* function in the R package phytools v. 0.6-99 (Revell, 2012). We used AICc to choose among different transition rate models (ER equal rates, SYM, symmetrical rates, and ARD all rates different).

## RESULTS

### Comparison of concatenation and species tree methods

Concatenated analyses based on nucleotides or proteins placed hemiodontids as sister to serrasalmids with strong support (99/98 and 94/93 UFBoot/SH-aLRT, respectively). In contrast, the MSC approach produced a polytomy among three families, Serrasalmidae, Hemiodontidae, and Cynodontidae, albeit with strong support for this trichotomy (PP >0.95). All other relationships among main characiform lineages are consistent with higher-level phylogenetic studies of the order (Betancur-R. et al., 2019; Supplements).

Concatenated and MSC approaches supported three major clades within Serrasalmidae (Fig. 2), with some disagreement on relationships within these clades. Namely, concatenated amino acid and MSC approaches support a different relationship among the three major lineages within the genus *Serrasalmus*, relative to the concatenated nucleotide analysis (Fig. 2). These three major lineages are denoted in Figure 2 as the ‘*rhombeus*’ clade (R), the short-snouted (brachycephalic) ‘*aureus*’ clade (A), and the ‘*maculatus*’ clade (M). Both the concatenated and MSC approaches applied to the amino acid dataset resolved the *Serrasalmus maculatus* clade (M) as sister to the remaining *Serrasalmus* taxa, the [‘*rhombeus*’clade (R) + *Pristobrycon calmoni*] and the ‘*aureus*’ clade (A), with high support, but this relationship differs in the concatenated nucleotide analysis (Fig. 2). All analyses resolve *Pygocentrus* as sister to *Serrasalmus* and the associated *Pristobrycon* species nested therein, with high support (Fig. 2). (Supplements)

**Figure 2.**
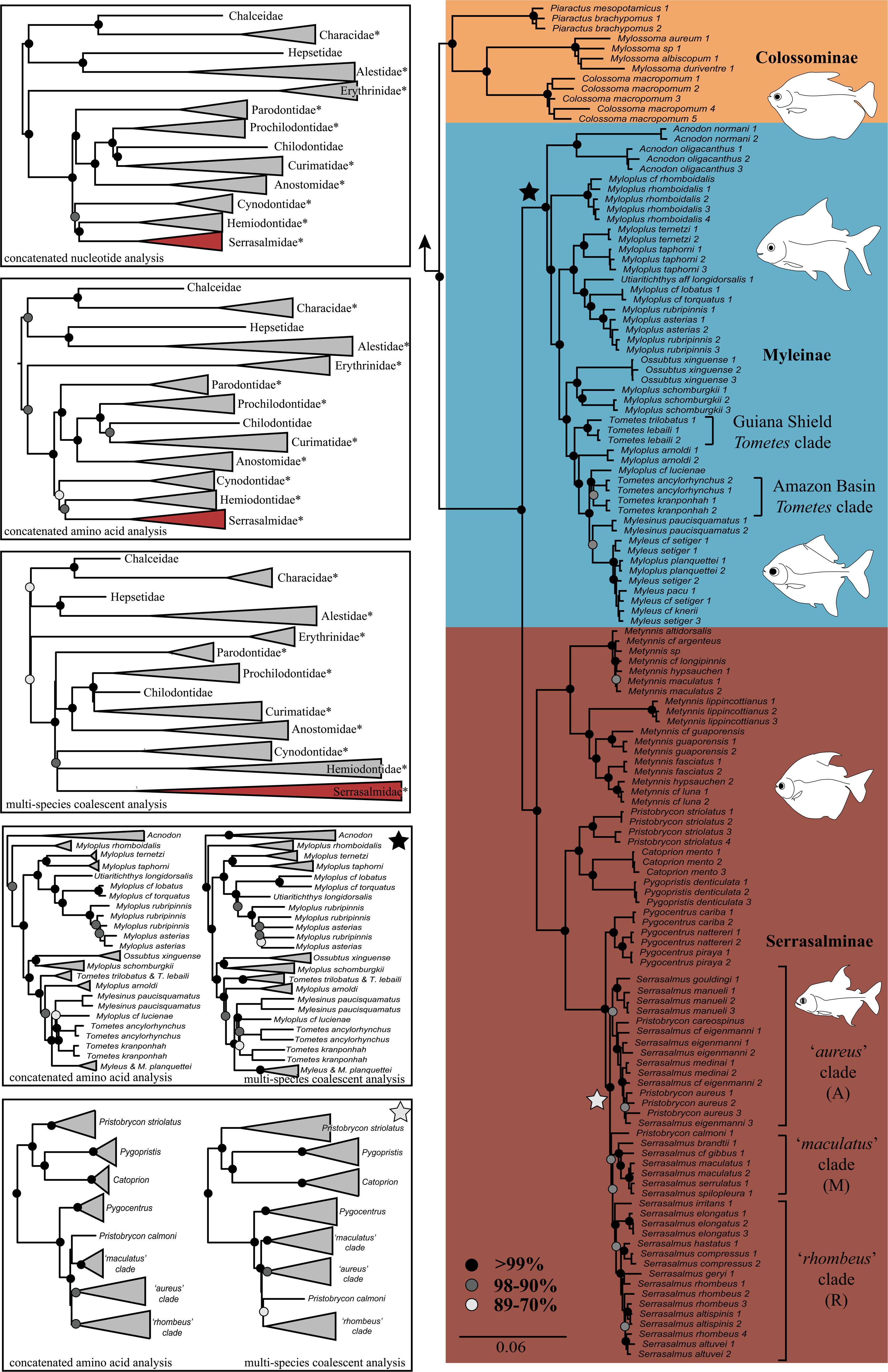
Comparison of concatenated nucleotide and amino acid trees and clade support in Serrasalmidae. Black circles represent nodes with > 99% UFBoot and aLRT support. Proposed morphological synapomorphies for Serrasalmidae and Colossominae discussed in Supplemental Materials.

The concatenated amino acid analysis and the coalescent analysis resolve *Mylesinus paucisquamatus* as sister to the clade comprised of *Tometes ancylorhynchus, T. kranponhah*, and *Myloplus lucienae*, albeit with low support in the amino acid dataset (Fig. 2). Alternatively, the concatenated nucleotide dataset resolves *Mylesinus paucisquamatus* as sister to the *Myleus* + *Myloplus planquettei* clade with moderately high support. The placement of *Utiaritichthys* differed between the concatenation and MSC analyses, though it was always nested within *Myloplus* (Fig. 2). For the first time, we can report that the newly described *Myloplus taphorni* (Andrade et al., 2019b), an endemic to the Mazaruni Basin in Guyana, is sister to *Myloplus ternetzi*, a northern Guiana Shield endemic. Our study also firmly resolves *Acnodon* as monophyletic and sister to all remaining members of Myleinae, something that all previous studies had found difficult to resolve with certainty (Fig. 2).

### Time calibration

Bayesian analyses in BEAST2 converged on estimates of the posterior distributions, as indicated by ESS values > 200 for all parameters. For each fossil calibration scheme, independent runs based on each of the four 50-gene subsets produced very similar mean-age estimates for all nodes. In contrast, we note marked differences between the two fossil calibration schemes with most obvious discrepancies in the estimated age for the MRCA of Serrasalmidae (HPD = 38-42.9 mya *vs*. 61-64.6 mya) and the split between serrasalmids and their hemiodontid sister clade (HPD = 55.5-76 mya *vs*. 68-81), although the latter estimates overlap considerably (Fig. 3). However, the ages of more inclusive clades overlap considerably between both dating schemes. Piranhas (Serrasalminae) are Miocene in age, in agreement with Thompson et al. (2014), with the modern radiation of *Serrasalmus* (and *Pristobrycon*) stemming from Messinian time periods (5.3-7.2 mya). The split between *Pygocentrus* and *Serrasalmus* being only slightly older, Tortonian-Messinian (combined HPD = 6-9 mya; Fig. 3). The split between piranhas and their more herbivorous cousins, *Metynnis* (both Serrasalminae), occurred on a long branch during the mid-Oligocene to mid-Miocene (combined HPD = 14-24 mya). While the radiation of carnivorous piranhas happened rapidly, the diversification of medium-sized pacus (Myleinae) was more gradual, occurring sometime within the same timeframe as *Metynnis* from piranhas (mid-Oligocene to mid-Miocene) (combined HPD = 11-30 mya; Fig. 3). Conversely, the largest pacu species (Colossominae) diverged from one another (*Piaractus* from *Mylossoma* + *Colossoma*) early on in their history, late Eocene to mid-Oligocene (combined HPD = 27-41 mya; Fig 3) according to Scheme 1 or earlier in Scheme 2, from late Paleocene to late Eocene (combined HPD = 39-63; Fig. 3).

**Figure 3.**
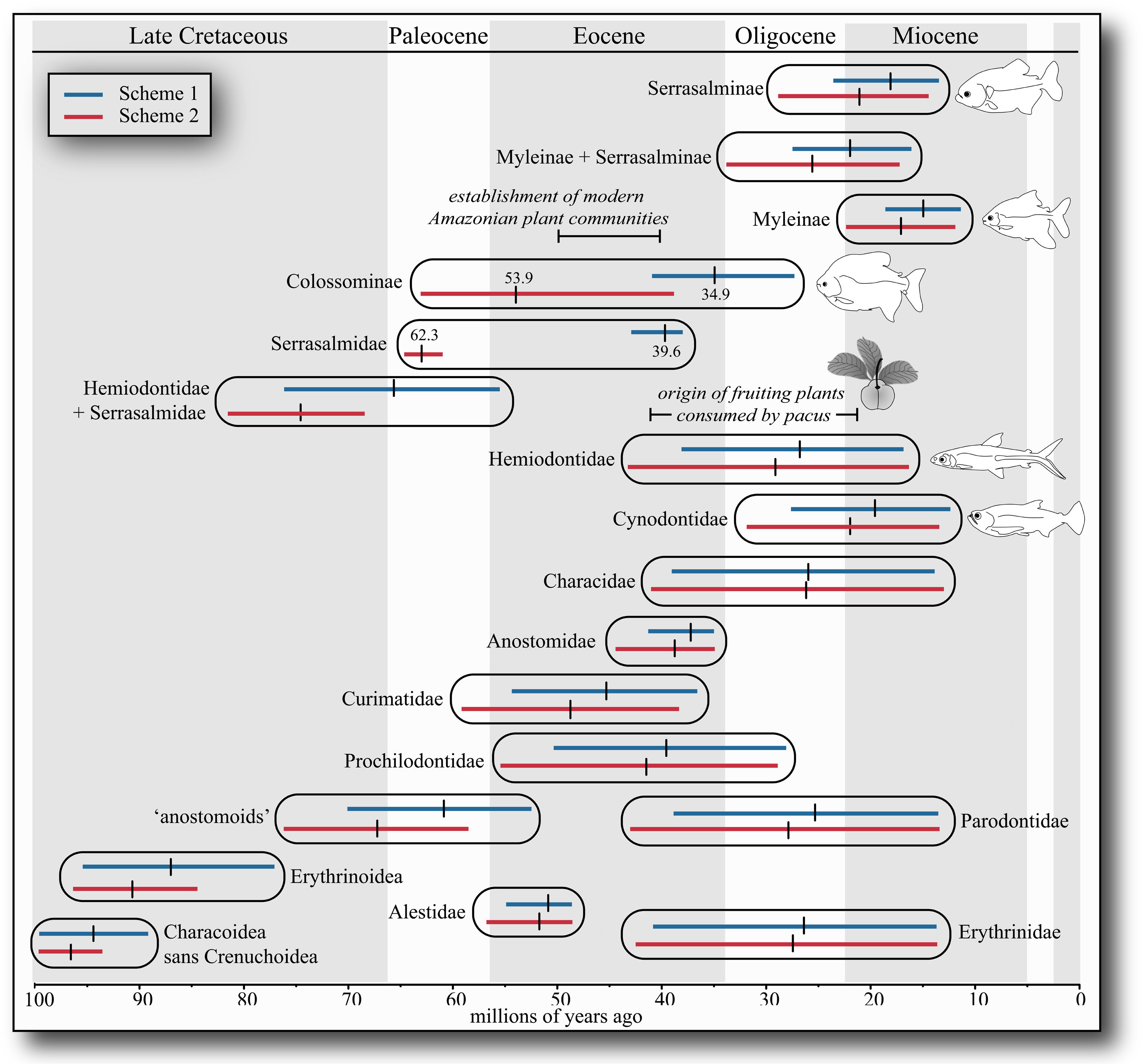
Comparison of HPD results for both time-calibration schemes. Blue bars represent HPD range from Scheme 1 and red bars represent HPD ranges from Scheme 2. Ranges are almost younger for Scheme 1, relative to Scheme 2. Raw mean estimated ages for the most divergent estimates are presented above their respective bars. Fruit icon is the rubber tree, *Hevea brasiliensis*, a preferred prey item of large-bodied pacus (Goulding, 1980).

### Diet reconstruction

For the more exclusive diet categories, the transition model with the lowest AICc (−102.5) was the ‘ARD’ or All Rates Different model, which allows transitions and reversions between every state to vary independently. Using this model, we found that shifts from piscivory or partial piscivorous feeding modes (i.e. fin- and scale-feeding) to omnivorous feeding modes where plants are consumed alongside flesh, are common in piranhas (Fig. 4). We also found equally probable support for obligate piscivory, omnivorous, or partial piscivorous ancestor for all piranhas. Strong support for obligate piscivory is inferred only at the nodes uniting *Pygocentrus* and the ‘*rhombeus*’ clade of *Serrasalmus*, respectively (Fig. 4). This is due to the generally omnivorous to outright herbivorous feeding habits of the sister taxa to all other piranhas, i.e. *Pygopristis denticulata* + *Pristobrycon striolatus*, which predominantly feed on plant materials as adults and their close relative, *Catoprion mento*, a scale-feeder (Nico & Taphorn, 1988; Nico & Morales, 1988).

**Figure 4.**
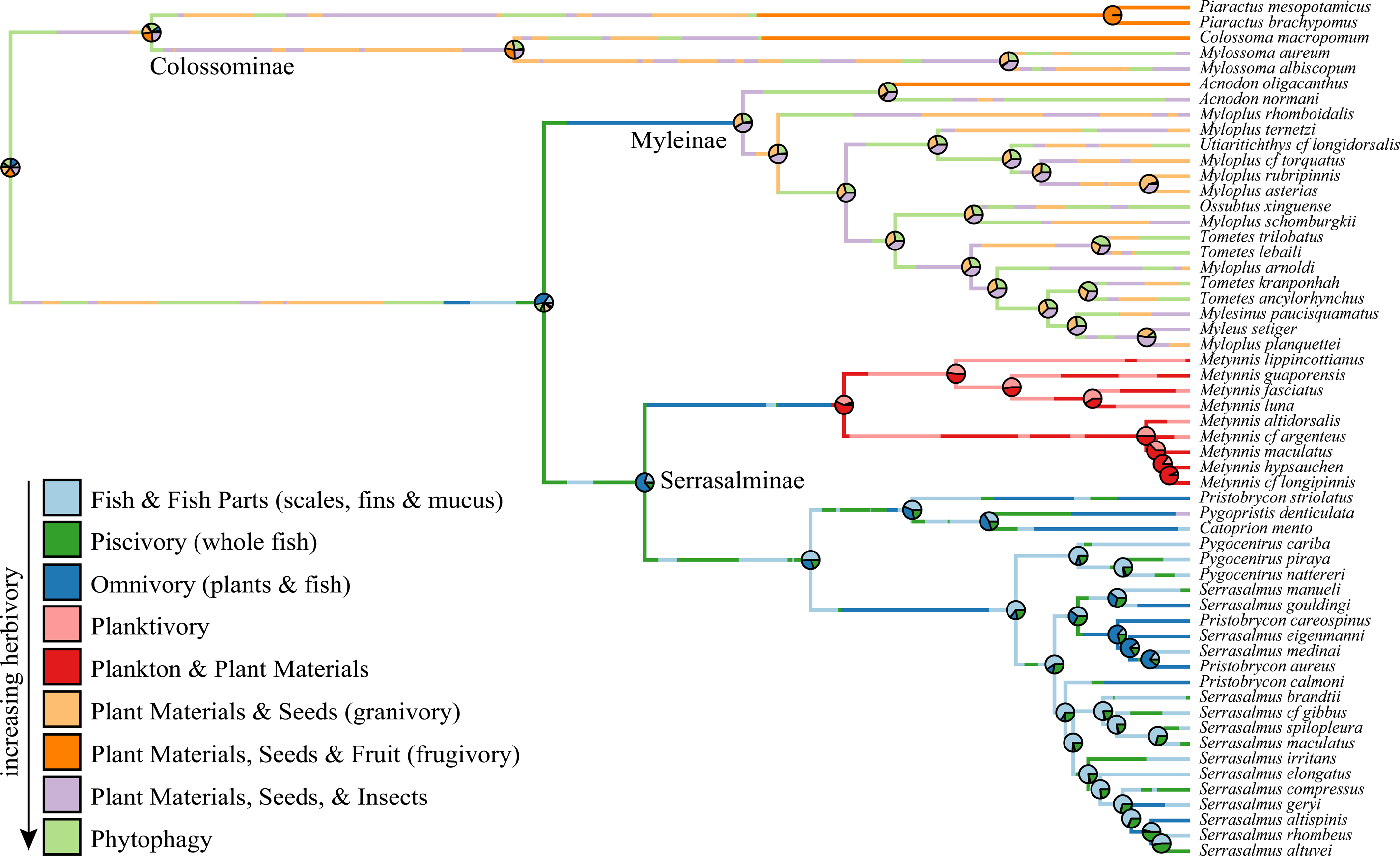
Discrete diet trait evolution in Serrasalmidae. (1) *FishFishParts*: Scales, fins, whole or partial fishes found in diet; (2) *Omnivory*: Whole or partial fishes & plants found in stomachs; (3) *Phytophagy*: fishes feeding either obligately or facultatively on riverweed (Podostemaceae); (4) *Piscivory*: feeding on whole or partial fishes; (5) *Planktivory*: obligate plankton feeders; (6) *Plankton & Plant Materials*: fishes feeding on plankton, algae, and other plant parts; (7) *Plant Materials & Seeds*: granivores feeding on seeds and other plant parts; (8) *Plant Materials, Seeds & Fruits*: frugivores, feeding on fruits, seeds, and other plant parts; (9) *Plant Materials, Seeds & Insects*: those species feeding on a combination of plant parts and invertebrates.

Also noteworthy is that reversals to more plant-based diets are characteristic of the brachycephalic ‘*aureus*’ (A) clade of piranhas, notably *Serrasalmus gouldingi* and the majority of *Pristobrycon* taxa, which feed as adults on fruits, seeds, leaves, insects and only occasional fish or fish parts (Nico & Taphorn, 1988; Prudente et al., 2016) (Fig. 4). Although several groups show increasing specialization for piscivory (e.g. *Pygocentrus*), this feeding mode does not appear to be an evolutionary dead-end for piranhas, with frequent transitions from piscivory to scale- and fin-feeding, or omnivorous and herbivorous diets (Supplements).

Our more comprehensive diet reconstructions document nuance among the diets of herbivorous taxa (Correa et al., 2007). Large-bodied pacus (*Colossoma, Piaractus*) are more frugivorous than smaller-bodied pacus. Also notable are the multiple instances of specialized ‘phytophagy’ in select taxa like *Tometes, Mylesinus, Ossubtus*, and *Utiaritichtys*, which specialize on the flowers, leaves, and stems of particular river weed plants found only in fast-flowing rapids (Pereira & Castro, 2014; Andrade et al., 2019) (Fig. 4). The paraphyly of *Tometes* in our phylogeny may suggest the sharp incisiform teeth critical in distinguishing *Tometes* from other myleines are misleading. Finally, several species of *Metynnis* are partially to entirely planktivorous, a novelty for serrasalmids and a hallmark for the *Metynnis* clade in general (Canan & Gurgel, 2002) (Fig. 4). However, feeding on plankton appears restrictive, with transitions out of obligate or facultative planktivory being exceedingly rare (Table 2; Fig. 4).

## DISCUSSION

### An inconvenient tooth: homoplasy, uncertainty, & taxonomic characters

From taxonomic identification to fossil calibrations, our understanding of serrasalmid evolution is substantially influenced by the selective regimes acting on tooth form and function. Authors like Géry (1977) used tooth morphotypes to distinguish between serrasalmid subfamilies and more recent studies have used these same tooth morphologies to make assumptions about diet (Huby et al., 2019), the latter classifying species as either herbivorous or carnivorous depending entirely on whether that taxon has either molariform or triangular teeth (respectively). Our findings reinforce the notion that serrasalmid diets are considerably more diverse than commonly believed, although these trends had been well-documented long ago (Goulding, 1980; Correa et al., 2007). Similarly, ‘herbivory’ and ‘carnivory’ are not ecological monoliths. For example, silver dollars or pacucitos (*Metynnis*) feed on plankton, whereas others medium-sized pacus consume insects (e.g. *Mylesinus*; Santos et al., 1997), or even scales (*Acnodon normani*; Leite & Jégu, 1990). Similarly, piranhas are not all piscivorous - many have reverted to more herbivorous feeding modes and some lineages (e.g. *Pygopristis, Pristobrycon striolatus*; Nico & Taphorn, 1988) are not ancestrally piscivorous. In fact, unlike other fishes, piscivory does not appear to limit niche evolution in piranhas, with closely related lineages displaying variable degrees of omnivory and piscivory, as well as ‘mutilating’ diet modes like fin- or scale-feeding (Collar et al., 2009; Collar et al., 2014; Fig. 4). Instead, the evolution of serrasalmid feeding structures and diets is far more diverse and complex than widely appreciated.

Tooth shape is not predictive of diet among most herbivorous and carnivorous piranhas. The specialized scale-feeding wimple piranha, *Catoprion mento*, has teeth unlike any of its relatives (Kolmann et al., 2018a, 2019), while *Catoprion*’s sister taxon *Pygopristis*, the only serrasalmid with pentacuspid teeth, feeds on plants and insects as an adult (Fink, 1989; Nico, 1991). But there are several notable cases of morphological convergence. Species assigned to the non-monophyletic genus *Tometes* share remarkably similar sharp, incisiform teeth, which we propose relates to convergence among species adapted to feeding on rheophilic river weed (Andrade et al., 2016; Huie et al., 2019). Similarly, all *Pristobrycon* species are noted for having short, deep skulls whereas most *Serrasalmus* have more elongate faces (Machado-Allison, 1985). Our data demonstrate that most short-snouted *Serrasalmus* (like *S. manueli* or *S. gouldingi*) and short-snouted *Pristobrycon* (‘*aureus*’ clade) are omnivorous. Shorter jaws are more effective at transferring jaw muscle forces to hard prey like seeds (Goulding, 1980), so dietary convergence among brachycephalic fishes may have misled morphological taxonomy again. These examples highlight the difficulty in translating diagnostic characters useful for identifying species in the field to phylogenetically informative characters that reflect shared evolutionary history. They also highlight how incorporating natural history observations (diet) and functional considerations can help taxonomists steer clear of homoplasy.

### Disagreement over the origin (time) of serrasalmids, but not their diversification timeline

We propose that a dating scheme which casts doubt on the taxonomic affinity of putative pacu teeth from Cretaceous deposits in Bolivia, is the more conservative approach to estimating the timeline of serrasalmid diversification. Isolated fossil material, particularly unarticulated teeth, are difficult to assign to specific taxonomic groups. Paleontologists studying heterodont elasmobranchs have a long-established history of skepticism when designating extinct species or evaluating taxonomic affinity based solely on isolated teeth (Shimada, 2005; Whitenack & Godfried, 2010; Marrama & Kriwet, 2017). Similarly, convergent and heterodont tooth morphologies are quite prevalent among serrasalmids, but also most characiforms in general (Murray et al., 2004a; Kolmann et al., 2019), and this phenomenon may lead taxonomic classification of fossils astray. Without a quantitative evaluation of tooth characters across pacus and relevant outgroups, we argue that assignment of isolated teeth to certain lineages or taxa is fraught with uncertainty.

The common practice of recycling fossil calibrations from study to study may lead to compounding issues with uncertainty and imprecision in our fossil calibration estimates. Although pacus have a well-documented fossil record (Lundberg et al., 1998, 2009; Gayet et al., 2001; Dahdul, 2007), the assumption that isolated teeth are the product of evolutionary stasis (Lundberg et al., 1986) may be premature without (1) broader consideration of outgroups, (2) evaluations of convergence across the phylogeny, or (3) discovery of more articulated skeletons. Given the notable teeth in serrasalmids, it is ironic that their sister taxon, unambiguously identified here as the hemiodontids, have miniscule teeth or are entirely edentulous (Roberts, 1971, 1974). At first glance, serrasalmids would appear to have more in common with their toothier distant relatives, the payaras (Cynodontidae) than hemiodontids; however, we note that our estimates for the divergence among these three clades is Paleocene-Eocene (55-81 mya),enough time for edentulism and near-edentulism to have evolved independently in both aardvarks (Afrotheria) and anteaters (Xenartha) (Upham et al., 2019).

The diversification of serrasalmids, particularly large-bodied, fruit-eating pacus, has been associated in the literature with the coincident diversification of fruiting plants (Correa et al., 2015). However, this shared co-evolutionary timeline between frugivorous fishes and their prey plants have primarily relied on stem ages of the asterids, rosids, and other plant groups like the spurges (Euphorbiaceae) (Horn et al., 2011; Correa et al., 2015). However, modern Amazonian plant communities are thought to have established themselves roughly ∼40-50 mya during the Eocene, and particularly those plants relying on ichthochory (fish-based seed dispersal; Jaramillo et al., 2010). Our proposed timeline for serrasalmid diversification (Fig. 3) is also Eocene (56-23 mya) in age, rather than Paleocene (66-56 mya), and corresponds better with the crown ages of particularly the most recent common ancestors of plant genera consumed by pacus today. *Colossoma macropomum* consumes fruits and seeds from rubber trees (*Hevea spruceana*), tucumã or jauari palms (*Astrocaryum* sp.), pouteria trees, and even the hallucinogenic iporuru plant (*Alchornea* sp.) (Goulding, 1980). Similarly, pirapitinga (*Piaractus*) also consume tucumã palm fruit, as well as fava (*Vicia faba*), and even luffa (Cucurbitaceae) (Goulding, 1980). The ages of these plant genera are all late Eocene to Oligocene in age (∼41-22 mya; Wojciechowski, 2003; Schaefer et al., 2009; Bartish et al., 2011; Roncal et al., 2012), intriguingly corresponding with the diversification timeline of frugivorous, large-bodied pacus (*Mylossoma, Colossoma*, and *Piaractus*) and most other serrasalmid genera. These estimates are also in line with the Eocene timeline for fruit-eating vertebrates like birds and mammals (Fleming & Kress, 2011).

### Suggestions for serrasalmid taxonomy, moving forward

The taxonomy and systematics of Serrasalmidae have a long-checkered past “fraught with confusion and instability” (Nico et al., 2018:172). Recent morphological studies have helped distinguish and diagnose a variety of valid genera and species (e.g., Pereira and Castro, 2014; Andrade et al., 2016a; 2016b; 2016c; 2017; 2018; 2019b; Ota et al., 2016; Mateussi et al., 2018; Nico et al., 2018; Escobar et al., 2019). The current study provides robust molecular support for recognizing three major lineages of Serrasalmidae at the subfamilial rank: Colossominae (pacus common to lowland, white water habitats), Myleinae (pacus common to upland clear- and black water habitats), and Serrasalminae (Metynnis and piranhas, cosmopolitan). Furthermore, there is strong support for the sister group relationship between Myleinae and Serrasalminae. Those results are consistent with previous phylogenies based on morphological (Cione et al., 2009) and molecular (Ortí et al., 2008; Thompson et al., 2014) data.

Based on our phylogenomic analysis of the family, we recommend the following (with more detail in the Supplements): (1) the establishment of a new subfamily delineating large-bodied pacus from myleine pacus, the Colossominae (see Supplements for proposed morphological synapomorphies); (2) that *Pristobrycon* Eigenmann 1915 should be synonymized (as has been repeatedly suggested) with *Serrasalmus* Lacepède 1803. Despite disagreement among our tree-building methods regarding the exact placement of *P. calmoni*, the type species for *Pristobrycon*, the taxon is repeatedly found nested within *Serrasalmus* (Fig. 2; Thompson et al., 2014). Moreover, the distinguishing characteristic of *P. calmoni*, the presence of a pre-anal serra, is shared with *Serrasalmus* but not with half of other *Pristobrycon* species (Machado-Allison, 2002). (3) As such, *Pristobrycon striolatus* (and the cryptic *P. scapularis*; Andrade et al., 2019) should also be elevated to its own generic rank. Finally, *Myloplus* should be broken into several genera, particularly *Myloplus rhomboidalis*, which was formerly known as *Prosomyleus*. Our phylogenetic framework can serve as a guide for future systematic reappraisals of the family, particularly at the genus-level, although the group still requires significant morphological reassessment going forward.

## Acknowledgments

We dedicate this manuscript to the late Javier Maldonado-Ocampo, an excellent companion when fishing for piranhas as well as a great colleague and friend. J Armbruster and D Werneke (AUM), NK Lujan (AMNH) and S Willis for field collections and tissues. M Zur, M Burridge, E Holm, and D Stacy (ROM). M Arce-H (ANSP) and R Singer and D. Nelson (UMMZ) for curatorial support of this work. D Taphorn (ROM), E Liverpool (U Guyana) assisted with specimen collection and identification. R Vullo for helpful advice regarding characiform fossils. M Girard and S Mueller for sending rare and yet surprisingly pertinent papers. R Peterson, H Saad, and V Rodriguez for help with DNA extraction and quantification. A Elbassiouny and O Lucanus for aid with specimens. HLF is grateful to M. Andrade for help identifying Guiana Shield specimens at the UMMZ. All analyses were performed on the Colonial One High Performance Computing System at George Washington University. MAK was supported by National Science Foundation (NSF) Post-Doctoral Fellowship in Biology (NSF-DBI 1712015). This study was funded by NSF-DEB-1541554 to G.O. and NSF-DEB-1929248 and NSF-DEB-1932759 to R.B.R. HLF was funded by Discovery Grant RGOIN-2014-05374 from the Natural Science and Engineering Research Council of Canada, the Royal Ontario Museum and the University of Michigan.

## SUPPLEMENTS

S1: Figure of previous evolutionary hypotheses among serrasalmid genera.

S2: Fossil dating & time calibration.

S3: Fossil dating & time calibration references.

S4: Serrasalmid diet table.

S5: Diet transition matrix from SIMMAP simulations.

S6: Serrasalmid diet references.

S7: Taxonomic recommendations & morphological synapomorphies.

S8: Table of museum specimens, tissue #s, cat #s, locale.

