## Supplemental Materials S1-S7 for "Phylogenomics of piranhas and pacus (Serrasalmidae) uncovers how convergent diets obfuscate traditional morphological taxonomy"

**S1 - PREVIOUS MORPHOLOGICAL & MOLECULAR PHLOGENETIC HYPOTHESES FOR SERRASALMIDAE**

(reproduced and modified from their original papers)

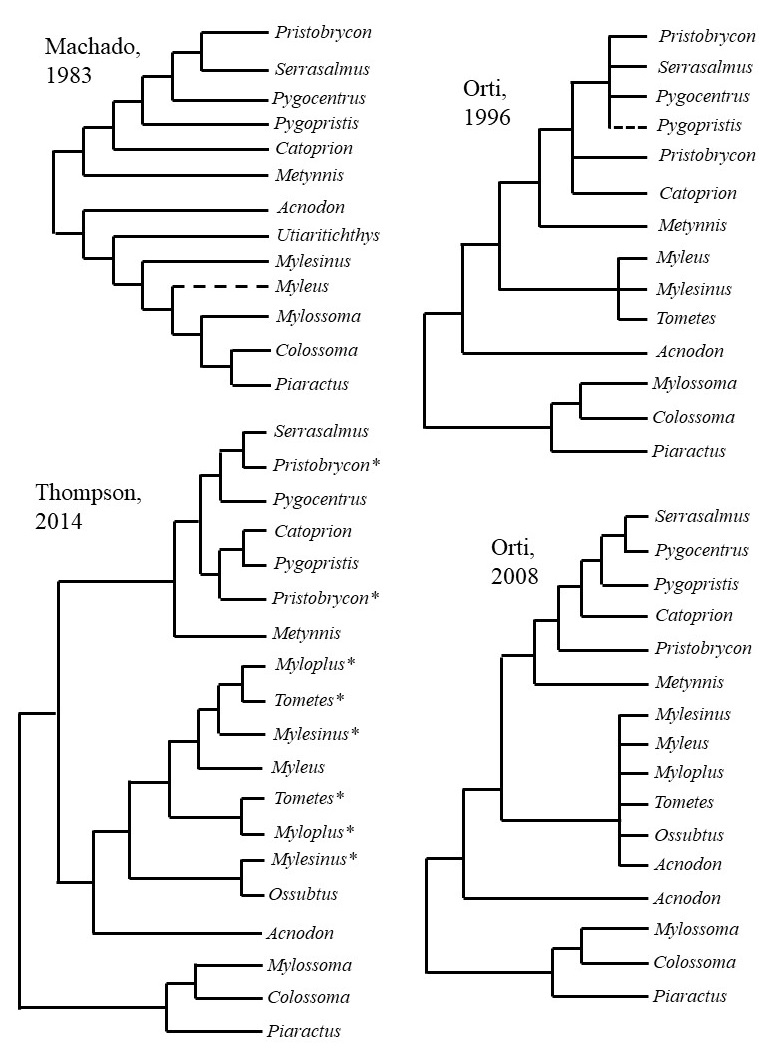

**S2 - DESCRIPTION & RATIONALE OF FOSSIL CALIBRATIONS**

We used BEAST 2 (v2.5.0; Bouckaert et al., 2014) to generate relaxed-clock divergence time estimates (Drummond et al., 2006) on four 50-gene subsets of our data, randomly selected from the 200 most complete genes, and pruned so that only one tip per taxon remained. First, we converted the concatenated nucleotide phylogeny estimated from IQTree (above) to a chronogram under penalized likelihood using the *chronos* function in R (ape v. 5.3; Paradis et al., 2019). This chronogram was used as a starting tree for the BEAST 2 analyses (Supplemental Files), the topology of the resulting trees was also constrained to match the concatenated nucleotide phylogeny. Each subset of 50 genes was run independently in BEAST 2 twice for 200,000,000 generations. All subsets had 103 included taxa, and Subset 1 had 4425 sites, Subset 2 had 5264 sites, Subset 3 had 5288 sites, and Subset 4 had 5096 sites. For each BEAST2 run, we used the GTR + gamma as our site model for each locus. We used a birth-death model tree prior for node time estimation, allowing for both speciation and extinction rates to vary for any given lineage (Drummond et al., 2006). We fixed the topology of our starting tree by turning off the following operators in BEAST2: (1) set ‘wide-exchange’ ="false", (2) set ‘narrow-exchange’ to “false”, (3) set ‘subtree-slide’ to 0, and (4) set ‘Wilson-Balding’ to 0.

To explore how the ambiguity surrounding these fossils alters our estimates of serrasalmid diversification, we used two different fossil calibration schemes and contrast the timelines produced by these analyses (and by previous studies, e.g. Broughton et al., 2013; Burns & Sidlauskas, 2019). We calibrated Scheme 1 with 15 fossil calibrations and Scheme 2 with 14 fossils. We used exponential distributions on each fossil prior except for the root, which used a normal distribution (Chen et al., 2010), in order to account for increasing uncertainty at further points in the past. Mean and standard deviations were estimated based on the calibration setting from other studies (e.g. Broughton et al., 2013; Chen et al., 2010; Thompson et al., 2014; Burns & Sidlauskas, 2019) which used the same fossils as calibrations points.

The first eleven fossil calibrations dealt with calibrations external to Serrasalmidae, in other characiform families. Within Characoidea, we dated the divergence between Characidae and Chalceidae, using fossil *Paleotetra* from the Aiuruoca Tertiary Basin (Weiss et al., 2012, 2014), Minas Gerais State in Eocene-Oligocene sediments (Garcia et al., 2000) (minimum age/offset = 23.0 mya, mean = XX). Two fossils were used to date within Alestoidea; for dating the base of Alestoidea *sans* Hepsetidae, we used fossil †*Alestoides eocaenicus* from Eocene Dormaal, near Brabant, Belgium (minimum age/offset = 48.6 mya, mean = 3.2) (Zanata & Vari, 2005; Gaudant & Smith, 2008; Chen et al., 2013). We also used fossils of the extant genus *Hydrocynus* to date the divergence between *Hydrocynus* + *Micralestes*, from the middle Eocene Hamada of Méridja deposits, in southwestern Algeria (Hammouda et al., 2016) (minimum age = 37.0 mya/offset, mean = 3.85).

We used two fossils pertaining to Erythrinidae; firstly, we used fossils attributed to Erythrinoidea (Gayet et al., 2003) from the Late Cretaceous to Paleocene of Bolivia (Gayet & Brito, 1989; Gayet, 1991; Gayet and Meunier, 1998) to date the root of our phylogeny, i.e. the node uniting Characoidea with Curimatoidea + Alestoidea (*sensu* Betancur et al., 2019) (minimum age/offset = 58.2 mya, mean = 13.82). To calibrate the node uniting *Hoplerythrinus + Hoplias*, we used teeth attributed to †*Paleohoplias assisbrasiliensis* (Gayet et al., 2003) from the late Miocene Solimões Formation of Acre State, Brazil (Latrubesse et al., 1997; Cione et al., 2003; Grosse et al., 2011) (minimum age = 7.2 mya/offset, mean = 17.0). Finally, we used fossil cynodontid teeth to calibrate the node uniting *Hydrolycus +* [*Rhaphiodon, Cynodon*]. These fossils are from middle Miocene sediments associated with the La Venta fauna near Tolima, Colombia (minimum age/offset = 7.2 mya, mean = 17.0; Lundberg, 1997; Cione & Casciotta, 2010).

Three fossils were used to date within anostomoids and one fossil from related Parodontidae; for dating the base of Anostomidae, we used a fossil oral tooth attributed to *Leporinus* sp. from the lower Pozo Formation, Contamana, Peru (Antoine et al. 2016), Middle Eocene sediments (minimum age/offset = 35.0 mya, mean = 7.7; Burns & Sidlauskas, 2019). We also used fossils of †*Leporinus scalabrinii* (Bogan et al., 2012) to date the divergence between *Abramites hypselonotus* + *Leporinus striatus*, from the late Miocene deposits of the Ituzaingó Formation in Entre Ríos, Argentina (Marshall et al., 1983; Cione et al., 2000, 2009) (minimum age/offset = 6 mya, mean = 9.7). Finally, to calibrate the node uniting *Cyphocharax + Psectrogaster* with *Curimata*, we used †*Cyphocharax mosesi* from the Tremembe Formation, Sao Paulo, Brazil in Oligocene sediments (Malabarba, 1996) (minimum age = 23.0 mya/offset, mean = 11.7). †*Cyphocharax mosesi* was originally proposed as forming a polytomy with the genera *Cyphocharax, Curimatella,* and *Steindachnerina* (Malabarba, 1996; Burns & Sidlauskas, 2019).

We also used fossil teeth attributed to *Parodon* by Roberts (1975) to date the divergence between *Apareiodon* + *Parodon*, from mid-late Miocene deposits of the Loyola Formation near La Cuenca, Ecuador (Bristow, 1973) (minimum age/offset = 11.2 mya, mean = 15.7) (Hungerbühler et al 2002).

Within Serrasalmidae, four fossil calibrations were used; firstly, for Scheme 2, we used the isolated pacu teeth first described in Gayet (1991), and used by Broughton et al. (2013) and Thompson et al. (2014) to date the divergence of serrasalmids from other non-serrasalmid characiforms (minimum age = 61.0 mya/offset, mean = 12.9). Whereas Broughton et al. used this fossil to represent the MRCA for *Pygocentrus* + *Hemiodus*, Thompson et al. used these fossil teeth to calibrate the node uniting *Serrasalmus* + *Piaractus*. For Scheme 1, we removed this calibration and replaced with pacu teeth described by DeCelles & Horton (2003) from the Paleocene-Eocene Santa Luca Formation, Bolivia (minimum age = 38.0 mya/offset, mean = 6.75). To calibrate the node uniting *Colossoma* + *Mylossoma*, we used teeth and partially articulated jaws documented by Lundberg et al. (1986) and Dahdul (2004) from the Miocene Castillo Formation, Venezuela (Rincon et al., 2014) (minimum age/offset = 17.2 mya, mean = 7.0). The pacu fossils from above predate fossils of *Piaractus* (Sanchez-Villagra & Aguilera, 2006) from the Tortonian Urumaco Formation in Falcón State, Venezuela (Dahdul, 2004). Next, we used fossil teeth attributed to indeterminate myleines (medium-sized pacus) to calibrate the MRCA of *Acnodon* + *Myloplus* (Roberts, 1975; Dahdul, 2004) from the mid-late Miocene Loyola Formation near La Cuenca, Ecuador (Bristow, 1973) (minimum age/offset = 11.2 mya, mean = 9.0) (Hungerbühler et al 2002; Dahdul, 2004). Finally, to calibrate the MRCA of all piranha genera, we used the upper Miocene fossil premaxilla described as †*Megapiranha paranensis* discovered in Entre Ríos, Argentina (Cione et al., 2009) (minimum age/offset = 6.8 mya, mean = 10.4).

Convergence of each gene subset was assessed individually in Tracer (v. 1.7.1) by checking that ESS values were greater than 200 for all parameters. Independent runs from each of the four different subsets were combined in LogCombiner if their 95% highest posterior densities for divergence times overlapped, and a maximum clade credibility tree was generated in TreeAnnotator for each of the two calibration schemes.

**S3 - TIME CALIBRATION, GEOLOGICAL DATING, & FOSSIL REFERENCES**

Betancur‐R, R., Arcila, D., Vari, R.P., Hughes, L.C., Oliveira, C., Sabaj, M.H. and Ortí, G., 2019. Phylogenomic incongruence, hypothesis testing, and taxonomic sampling: The monophyly of characiform fishes. Evolution, 73(2), pp.329-345.

Bogan, S., Sidlauskas, B., Vari, R.P. and Agnolin, F., 2012. Arrhinolemur scalabrinii Ameghino, 1898, of the late Miocene: a taxonomic journey from the Mammalia to the Anostomidae (Ostariophysi: Characiformes). Neotropical Ichthyology, 10(3), pp.555-560.

Bristow, C.R. 1973. Guide to the geology of the Cuenca Basin, southern Ecuador. Ecuadorian Geological and Geophysical Society.

Broughton, R.E., Betancur-R, R., Li, C., Arratia, G. and Ortí, G., 2013. Multi-locus phylogenetic analysis reveals the pattern and tempo of bony fish evolution. PLoS currents, 5.

Burns, M.D. and Sidlauskas, B.L., 2019. Ancient and contingent body shape diversification in a hyperdiverse continental fish radiation. Evolution, 73(3), pp.569-587.

Cione, A. L., M. M. Azpelicueta, M. Bond, A. A. Carlini, J. R. Casciotta, M. A. Cozzuol, M. de la Fuente, Z. Gasparini, F. J. Goin, J. Noriega, G. J. Scillato-Yané, L. Soibelzon, E. P. Tonni, D. Verzi, & M. G. Vucetich. 2000. Miocene vertebrates from Entre Ríos Province, eastern Argentina. Pp. 191-237. In: Aceñolaza, F. G. & R. Herbst (Eds.). El Neógeno de Argentina. INSUGEO Serie Correlación Geológica, 14.

Cione, A.L., Dahdul, W.M., Lundberg, J.G. and Machado-Allison, A., 2009. *Megapiranha paranensis*, a new genus and species of Serrasalmidae (Characiformes, Teleostei) from the upper Miocene of Argentina. Journal of Vertebrate Paleontology, 29(2), pp.350-358.

Cione, A.L. and Casciotta, J.R., 1997. Miocene cynodontids (Osteichthyes: Characiformes) from Paraná, central eastern Argentina. Journal of Vertebrate Paleontology, 17(3), pp.616-619.

Dahdul, W.M., 2004. Fossil serrasalmine fishes (Teleostei: Characiformes) from the Lower Miocene of north-western Venezuela. Fossils of the Miocene Castillo Formation, Venezuela: contributions on neotropical palaeontology, (71), pp.23-28.

Dahdul, W.M., 2007. Phylogenetics and diversification of the neotropical Serrasalminae (Ostariophysi: Characiformes).

Dahdul, W.M., 2010. Review of the phylogenetic relationships and fossil record of Characiformes. Gonorynchiformes and ostriophysan relationships: A comprehensive review, pp.441-464.

DeCelles, P.G. and Horton, B.K. 2003. Early to middle Tertiary foreland basin development and the history of Andean crustal shortening in Bolivia. Geological Society of America Bulletin, 115(1), pp.58-77.

Gayet, M. and Brito, P.M., 1989. Ichtyofaune nouvelle du Crétacé supérieur du groupe Bauru (états de Sao Paulo et Minas Gerais, Brésil). Geobios, 22(6), pp.841-847.Gayet, M. and Meunier, F.J., 1998. Maastrichtian to early late Paleocene freshwater Osteichthyes of Bolivia: additions and comments. Phylogeny and classification of Neotropical fishes, pp.85-110.

Gayet, M., Meunier, F.J. 1998. Maastrichtian to early late Paleocene freshwater Osteichthyes of Bolivia: additions and comments. In: Malabarba, L.R., Reis, R.E., Vari, R.P., Lucena, Z.M., Lucena, C.A. (Eds.), Phylogeny and Classification of Neotropical Fishes. Edipucrs, Porto Alegre.

Gayet, M. 1991. Holostean and Teleostean fish from Bolivia. R. Suarez-Soruco (ed.), Fosiles y Facies de Bolivia, I. Revista Tehcnica de YPFB 12; pp. 453–494.

Gayet, M., Marshall, L.G., Sempere, T., Meunier, F.J., Cappetta, H. and Rage, J.C., 2001. Middle Maastrichtian vertebrates (fishes, amphibians, dinosaurs and other reptiles, mammals) from Pajcha Pata (Bolivia). Biostratigraphic, palaeoecologic and palaeobiogeographic implications. Palaeogeography, Palaeoclimatology, Palaeoecology, 169(1-2), pp.39-68.

Gayet, M., Jégu, M., Bocquentin, J. and Negri, F.R., 2003. New characoids from the Upper Cretaceous and Paleocene of Bolivia and the Mio-Pliocene of Brazil: phylogenetic position and paleobiogeographic implications. Journal of Vertebrate Paleontology, 23(1), pp.28-46.

Garcia, M. J., Santos,M. & Hasui, Y. 2000. Palinologia da parte aflorante da Formac¸ ˜ao Entre-C´orregos, Bacia de Aiuruoca, Terci´ario do Estado de Minas Gerais, Brasil. Revista Universidade de Guarulhos, 5, 259.

Gross, M., Piller, W.E., Ramos, M.I. and da Silva Paz, J.D., 2011. Late Miocene sedimentary environments in south-western Amazonia (Solimões formation; Brazil). Journal of South American Earth Sciences, 32(2), pp.169-181.

Hungerbühler, D., Steinmann, M., Winkler, W., Seward, D., Egüez, A., Peterson, DE, Helg, U. and Hammer, C., 2002. Neogene stratigraphy and Andean geodynamics of southern Ecuador. Earth-Science Reviews, 57 (1-2), pp. 75-124.

Latrubesse E.M., Bocquentin J., Santos J.C.R., Ramonell C.G. 1997. Paleoenvironmental model for the late Cenozoic of southwestern Amazonia: paleontology and geology. Acta Amazonica. 27:103–118.

Lundberg, J.G., Machado-Allison, A. and Kay, R.F., 1986. Miocene characid fishes from Colombia: evolutionary stasis and extirpation. Science, 234(4773), pp.208-209.

Lundberg, J.G., 1998. The temporal context for the diversification of Neotropical fishes. Phylogeny and classification of Neotropical fishes, pp.49-68.

Lundberg, J.G., Sabaj Pérez, M.H., Dahdul, W.M. and Aguilera, O.A., 2009. The Amazonian neogene fish fauna. Amazonia: Landscape and Species Evolution: A look into the past, pp.281-301.

Malabarba, M. 1996. Reassessment and relationships of *Curimata mosesi* Travassos & Santos, a fossil fish (Teleostei: Characiformes: Curimatidae) from the tertiary of São Paulo, Brazil. Comunicações do Museu de Ciências da PUCRS, Série Zoologia 9:55-63.

Marshall, L., R. Hoffstetter & R. Pascual. 1983. Mammals and stratigraphy: geochronology of the continental mammal-bearing Tertiary of South America. Palaeovertebrata Mémoire Extraordinaire, 1-93.

Otero, O., Valentin, X. and Garcia, G., 2008. Cretaceous characiform fishes (Teleostei: Ostariophysi) from Northern Tethys: description of new material from the Maastrichtian of Provence (Southern France) and palaeobiogeographical implications. Geological Society, London, Special Publications, 295(1), pp.155-164.

Patterson, C. 1993. Osteichthyes: Teleostei. In: Benton, M.J. (ed.) The Fossil Record 2, 622-656.

Roberts, T.R. 1975. Characoid fish teeth from the Miocene deposits in the Cuenca Basin, Ecuador. Journal of Zoology. 175(2): 259-271.

Sánchez‐Villagra, M.R. and Aguilera, O.A., 2006. Neogene vertebrates from Urumaco, Falcón State, Venezuela: diversity and significance. Journal of Systematic Palaeontology, 4(3), pp.213-220.

**S4 - TAXON & DIET MATCHING TABLE**

| **Species** | **Diet Category** | **Reference(s)** |
| --- | --- | --- |
| *Acnodon normani* | PlantMatSeedsInverts | Leite & Jégu, 1990 |
| *Acnodon oligacanthus* | PlantMatSeedsFruit | Planquette et al., 1996; Mol, 2012 |
| *Catoprion mento* | FishFishParts | Vieira & Gery, 1979; Nico & Taphorn, 1988; Nico & Morales, 1994; Wantzen et al., 2002 |
| *Colossoma macropomum* | PlantMatSeedsFruit | Goulding, 1980; Goulding & Carvalho, 1982; Lucas, 2008 |
| *Metynnis altidorsalis* | PlanktonPlantMat | Mol, 2012; do Carmo, 2013; Ota, 2015 |
| *Metynnis argenteus* | Planktivory | do Carmo, 2013 |
| *Metynnis fasciatus* | Planktivory | Ota, 2015 |
| *Metynnis guaporensis* | PlanktonPlantMat | Andrade et al., 2019; Ota, 2015 |
| *Metynnis hypsauchen* | PlanktonPlantMat | Araujo-Lima et al., 1986; Ota, 2015 |
| *Metynnis lippincottianus* | PlanktonPlantMat | Canan & Gurgel 2002 (as *M. roosevelti*); Ramos et al., 2008 |
| *Metynnis longipinnis* | PlanktonPlantMat | Zarske & Géry, 2008; Ota, 2015 |
| *Metynnis luna* | Planktivory | Ota, 2015; Andrade et al., 2019 |
| *Metynnis maculatus* | PlanktonPlantMat | Silva-Camacho et al., 2014; Pelicice & Agostinho, 2006 |
| *Mylesinus paucisquamatus* | PlantMatSeedsInverts | Santos et al., 1997; Dary et al., 2017 |
| *Myleus setiger* | PlantMatSeedsInverts | Dary et al., 2017; Andrade et al., 2019 |
| *Myloplus arnoldi* | PlantMatSeeds | Zuluaga-Gómez et al. 2016 |
| *Myloplus asterias* | PlantMatSeeds | Nico, 1991; Dary et al 2017; Andrade et al., 2019 |
| *Myloplus planquettei* | PlantMatSeeds | Jegu et al., 2003 |
| *Myloplus rhomboidalis* | PlantMatSeedsInverts | Boujard et al., 1990; Andrade et al., 2019 |
| *Myloplus rubripinnis* | PlantMatSeeds | Dary et al., 2017; Gonzalez & Vispo, 2002; Andrade et al., 2019 |
| *Myloplus schomburgkii* | PlantMatSeedsInverts | Zuluaga-Gómez et al., 2016; Dary et al., 2017; Andrade et al., 2019 |
| *Myloplus torquatus* | PlantMatSeeds | Nico, 1991; Dary et al., 2017 |
| *Mylossoma aureum* | PlantMatSeedsInverts | Soares et al. 1986; Dos Santos 1990; Pouilly et al., 2003, 2004 |
| *Mylossoma albiscopum* | PlantMatSeedsInverts | Gonzalez & Vispo, 2003; Pouilly et al., 2003, 2004 |
| *Ossubtus xinguense* | Phytophagy | Jegu, 1992; Andrade et al., 2016b |
| *Piaractus brachypomus* | PlantMatSeedsFruit | Goulding, 1980; Lucas, 2008 |
| *Piaractus mesopotamicus* | PlantMatSeedsFruit | Galetti et al., 2008; Sório et al., 2014 |
| *Pristobrycon calmoni* | Omnivory | Gonzalez & Vispo, 2002 (as *Pristobrycon* spp); Nico, 1991 |
| *Pristobrycon careospinus* | Omnivory | Gonzalez & Vispo, 2002 (as *Pristobrycon* spp); Nico, 1991 |
| *Pristobrycon striolatus* | Omnivory | Goulding, 1988; Nico & Taphorn, 1988; Nico, 1991 |
| *Pygocentrus cariba* | FishFishParts | Gonzalez & Vispo, 2002; Nico & Taphorn, 1988; Winemiller, 1989 |
| *Pygocentrus nattereri* | FishFishParts | Nico & Taphorn, 1988; Ferreira et al 2014 |
| *Pygocentrus piraya* | FishFishParts | Trindade & Jucá-Chagas, 2008 |
| *Pygopristis denticulata* | PlantMatSeedsInverts | Nico, 1991 (juveniles); Nico & Taphorn, 1998 |
| *Serrasalmus altispinis* | Omnivory | Andrade et al., 2019; dubious ID |
| *Serrasalmus altuvei* | Piscivory | Nico, 1991; Nico & Taphorn, 1998 |
| *Serrasalmus brandtii* | Piscivory | Pompeu, 1999; Gurgel et al., 2002; Trindade & Jucá-Chagas, 2008 |
| *Serrasalmus compressus* | Piscivory | Pouilly et al., 2003, 2004 |
| *Serrasalmus eigenmanni* | Omnivory | Pouilly et al 2003; dubious ID |
| *Serrasalmus elongatus* | FishFishParts | Nico & Taphorn, 1998; Röpke et al., 2014 |
| *Serrasalmus geryi* | FishFishParts | Araujo-Lima et al., 1995; do Carmo, 2013 |
| *Serrasalmus gouldingi* | Omnivory | Prudente et al., 2016 |
| *Serrasalmus eigenmanni* | Omnivory | Nico, 1991 |
| *Serrasalmus irritans* | FishFishParts | Nico & Taphorn, 1988 |
| *Serrasalmus maculatus* | Piscivory | Carvalho et al., 2007; Behr & Signor, 2008 |
| *Serrasalmus manueli* | Piscivory | Dary et al 2017; Nico, 1991 |
| *Serrasalmus medinai* | FishFishParts | Winemiller, 1989; Nico, 1991 |
| *Serrasalmus rhombeus* | FishFishParts | Nico & Taphorn, 1988; Gonzalez & Vispo, 2002; Pouilly et al., 2003 |
| *Serrasalmus spilopleura* | FishFishParts | Wantzen et al., 2002; Raposo & Gurgel, 2003 |
| *Tometes ancylorhynchus* | Phytophagy | Andrade et al., 2016, 2019 |
| *Tometes kranponhah* | Phytophagy | Andrade et al., 2016, 2018, 2019 |
| *Tometes lebaili* | Phytophagy | Mol, 2012; Jégu et al., 2002a |
| *Tometes trilobatus* | Phytophagy | Mol, 2012; Jégu et al., 2002b |
| *Utiaritichthys aff longidorsalis* | Phytophagy | Jegu etal., 1989; Pereira & Castro, 2014 |

**S5 - DIET STATE TRANSITION MATRIX**

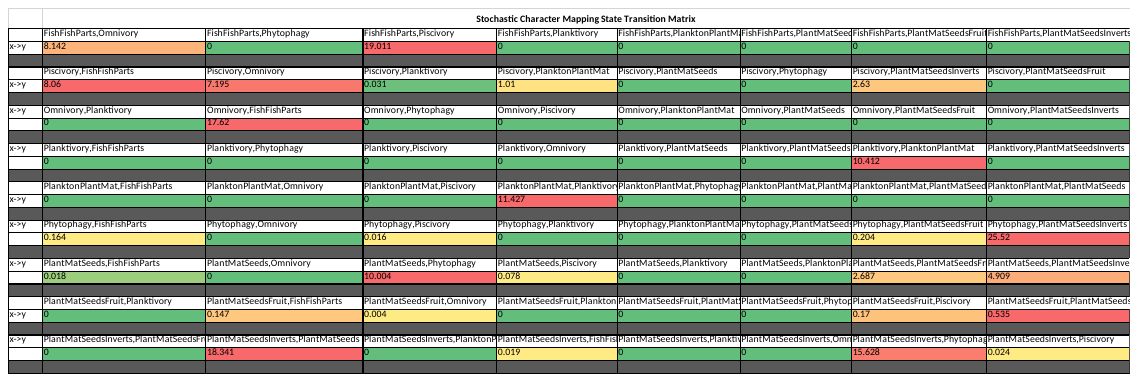

**S6 - DIET REFERENCES**

Anderson, J.T., Rojas, J.S. and Flecker, A.S., 2009. High-quality seed dispersal by fruit-eating fishes in Amazonian floodplain habitats. Oecologia, 161(2), pp.279-290.

Andrade, M. C., M. Jégu, and T. Giarrizzo. 2016a. *Tometes kranponhah* and *Tometes ancylorhynchus* (Characiformes: Serrasalmidae), two new phytophagous serrasalmids, and the first *Tometes* species described from the Brazilian Shield. Journal of Fish Biology 89(1): 467-94.

Andrade, M. C., L. M. Sousa, R. P. Ota, M. Jégu, and T. Giarrizzo. 2016b. Redescription and geographical distribution of the endangered fish *Ossubtus xinguense* Jégu 1992 (Characiformes, Serrasalmidae) with comments on conservation of the rheophilic fauna of the Xingu River. PLoS ONE 11(9): e0161398.

Andrade, M. C., V. N. Machado, M. Jégu, I. P. Farias, and T. Giarrizzo. 2017. A new species of *Tometes* Valenciennes 1850 (Characiformes: Serrasalmidae) from Tocantins-Araguaia River Basin based on integrative analysis of molecular and morphological data. PLoS ONE 12(4): e0170053.

Andrade, M. C., D. B. Fitzgerald, K. O. Winemiller, P. S. Barbosa, and T. Giarrizzo. 2018. Trophic niche segregation among herbivorous serrasalmids from rapids of the lower Xingu River, Brazilian Amazon. Hydrobiologia: 1-16.

Andrade, M.C., Winemiller, K.O., Barbosa, P.S., Fortunati, A., Chelazzi, D., Cincinelli, A. and Giarrizzo, T., 2019. First account of plastic pollution impacting freshwater fishes in the Amazon: Ingestion of plastic debris by piranhas and other serrasalmids with diverse feeding habits. Environmental Pollution, 244, pp.766-773.

Araujo‐Lima, C.A.R.M., Portugal, L.P.S. and Ferreira, E.G., 1986. Fish–macrophyte relationship in the Anavilhanas archipelago, a black water system in the Central Amazon. Journal of Fish Biology, 29(1), pp.1-11.

Araujo‐Lima, Carlos Alberto Rego Monteiro; Agostinho, Angelo Antonio; Fabre, Nidia N. Trophic aspects of fish communities in Brazilian rivers and reservoirs. In: Tundisi, José Galizia; Bicudo, Carlos Eduardo de Mattos; Matsumura- Tundisi, Takako (Ed.). Limnology in Brazil. Rio de Janeiro: ABC/SBL, 1995. p.[105]-136.

Behr, E.R. and Signor, C.A., 2008. Distribuição e alimentação de duas espécies simpátricas de piranhas *Serrasalmus maculatus* e *Pygocentrus nattereri* (Characidae, Serrasalminae) do rio Ibicuí, Rio Grande do Sul, Brasil. Iheringia. Série Zoologia, 98(4), pp.501-507.

Boujard, T., D. Sabatier, R. Rojas-Beltran, M.-F. Prevost, and J.-F. Renno. 2009. The food habits of three allochthonous feeding characoids in French Guiana. Revue d’écologie 45: 247-58.

Canan, B. and Gurgel, H.D.C.B. 2002. Feeding and diet rhythms of Metynnis roosevelti Eigenmann (Characidae, Myleinae) at Jiqui Lake, Parnamirim, Rio Grande do Norte, Brasil. Revista Brasileira de Zoologia. 19(2): 309-316.

Carvalho, L.N., Arruda, R., Raizer, J. and Del-Claro, K., 2007. Feeding habits and habitat use of three sympatric piranha species in the Pantanal wetland of Brazil. Ichthyological exploration of freshwaters, 18(2), p.109.

Correa, S.B., Winemiller, K.O., Lopez-Fernandez, H. and Galetti, M., 2007. Evolutionary perspectives on seed consumption and dispersal by fishes. Bioscience, 57(9), pp.748-756.

Correa, S.B. and Winemiller, K.O., 2014. Niche partitioning among frugivorous fishes in response to fluctuating resources in the Amazonian floodplain forest. Ecology, 95(1), pp.210-224.

Correa, S.B., Costa‐Pereira, R., Fleming, T., Goulding, M. and Anderson, J.T., 2015. Neotropical fish–fruit interactions: eco‐evolutionary dynamics and conservation. Biological Reviews, 90(4), pp.1263-1278.

Correa, S.B., Arujo, J.K., Penha, J., Nunes da Cunha, C., Bobier, K.E. and Anderson, J.T., 2016. Stability and generalization in seed dispersal networks: a case study of frugivorous fish in Neotropical wetlands. Proceedings of the Royal Society B: Biological Sciences, 283(1837), p.20161267.

Correa, S.B., de Oliveira, P.C., Nunes da Cunha, C., Penha, J. and Anderson, J.T., 2018. Water and fish select for fleshy fruits in tropical wetland forests. Biotropica. 50(2): 312-318.

Dary, E.P., Ferreira, E., Zuanon, J. and Röpke, C.P., 2017. Diet and trophic structure of the fish assemblage in the mid-course of the Teles Pires River, Tapajós River basin, Brazil. Neotropical Ichthyology, 15(4).

Da Silva, ATD, Zina, J, Ferreira, FC, Gomiero, LM, Goitein, R. (2015). Caudal fin-nipping by *Serrasalmus maculatus* (Characiformes: Serrasalmidae) in a small water reservoir: seasonal variation and prey selection. Zoologia Curitiba, 32, 457-462.

Do Carmo, C.M., 2013. Ecomorfologia e alimentação de peixes na bacia do Rio Das Mortes. (Doctoral dissertation, Universidade do Estado de Mato Grosso).

Ferreira, F.S., Vicentin, W., Costa, F.E.D.S. and Súarez, Y.R., 2014. Trophic ecology of two piranha species, *Pygocentrus nattereri* and *Serrasalmus marginatus* (Characiformes, Characidae), in the floodplain of the Negro River, Pantanal. Acta Limnologica Brasiliensia, 26(4), pp.381-391.

Galetti, M., Donatti, C.I., Pizo, M.A. and Giacomini, H.C., 2008. Big fish are the best: seed dispersal of *Bactris glaucescens* by the pacu fish (*Piaractus mesopotamicus*) in the Pantanal, Brazil. Biotropica, 40(3), pp.386-389.

Géry, J., 1977. Characoids of the World. T.F.H. Publications Inc., Neptune City, New Jersey.

González, N. & Vispo, C. 2002. Aspects of the diet and feeding ecologies of fish from nine floodplain lakes of the lower Caura, Venezuelan Guayana. Scientia Guaianae. 12: 329-3.

Goulding, M. 1980. The fishes and the forest: explorations in Amazonian natural history. University of California Press, Berkeley.

Goulding, M. & M. L. Carvalho, 1982. Life history and management of the tambaqui (*Colossoma macropomum*, Characidae): an important Amazonian food fish. Revista Brasileira de Zoologia 1: 107–133.

Gurgel, H.D.C.B., Lucas, F.D. and Souza, L.D.L.G., 2002. Dieta de sete espécies de peixes do semi-árido do Rio Grande do Norte, Brasil. Revista de Ictiologia, 10(1/2), pp.7-16.

Jégu, M., G. M. dos Santos, and E. J. Gondim Ferreira. 1989. Une nouvelle espèce du genre *Mylesinus* (Pisces, Serrasalmidae), *M. paraschomburgkii*, décrite des bassins du Trombetas et du Uatumã (Brésil, Amazonie). Revue d’Hydrobiologie Tropicale 22(1): 49-62.

Jégu, M. 1992. *Ossubtus xinguense*, nouveaux genre et espèce du Rio Xingu, Amazonie, Brésil (Teleostei: Serrasalmidae). *Ichthyological Exploration of Freshwaters*, 3, 235-252.

Jégu, M., P. Keith, and E. Belmont-Jégu. 2002a. Une nouvelle espéce de *Tometes* (Teleostei: Characidae: Serrasalminae) du bouclier Guyanais, *Tometes lebaili* n. sp. Bulletin Français de la Pêche et de la Pisciculture (364): 23-48.

Jégu, M., Santos, G.D., Keith, P. and Le Bail, P.Y., 2002b. Description complémentaire et réhabilitation de *Tometes trilobatus* Valenciennes, 1850, espèce-type de Tometes Valenciennes (Characidae: Serrasalminae). Cybium, 26(2), pp.99-122.

Jégu, M., P. Keith, and P.-Y. Le Bail. 2003. *Myloplus planquettei* sp. n. (Teleostei, Characidae), une nouvelle espèce de grand Serrasalminae phytophage du bouclier guyanais. Revue suisse de zoologie. 110: 833-53.

Leite, RG, Jégu, M. (1990). Régime alimentaire de deux espèces d'*Acnodon* (Characiformes, Serrasalmidae) et habitudes lepidophages de *A. normani*. *Cybium*, 14, 353-359.

Lucas, C.M. 2008. Within flood season variation in fruit consumption and seed dispersal by two characin fishes of the Amazon. Biotropica 40(5): 581-89.

Machado-Allison, A., and C. Garcia. 1986. Food habits and morphological changes during ontogeny in three serrasalmin fish species of the Venezuelan floodplains. Copeia 1986(1): 193.

Mol, J. H. 2012. The freshwater fishes of Suriname. Leiden, Brill Academic Publishers, 890 pp.

Nico, LG, Taphorn, DC. (1988). Food habits of piranhas in the low llanos of Venezuela. *Biotropica*, 311-321.

Nico, L. G. 1991. Trophic ecology of piranhas (Characidae: Serrasalminae) from savanna and forest regions in the Orinoco River basin of Venezuela. Ph.D. dissertation, University of Florida, Gainesville, pp 209.

Nico, L.G. and de Morales, M., 1994. Nutrient content of piranha (Characidae, Serrasalminae) prey items. Copeia, 1994(2), pp.524-528.

Northcote, TG, Northcote, RG, Arcifa, MS. (1986). Differential cropping of the caudal fin lobes of prey fishes by the piranha, *Serrasalmus spilopleura*. *Hydrobiologia*, 141, 199-205.

Ota, R.P., 2015. Revisão taxonômica e filogenia morfológica de Metynnis Cope, 1878 (Characiformes: Serrasalmidae).

Pelicice, F.M., Agostinho, A.A. and Thomaz, S.M., 2005. Fish assemblages associated with Egeria in a tropical reservoir: investigating the effects of plant biomass and diel period. Acta Oecologica, 27(1), pp.9-16.

Pereira, TN, Castro, R. (2014). A new species of *Utiaritichthys* Miranda Ribeiro (Characiformes: Serrasalmidae) from the Serra dos Parecis, Tapajós drainage. *Neotropical Ichthyology*, 12, 397-402.

Planquette, P., Keith, P. and Le Bail, P.Y., 1996. Atlas des Poissons dEau Douce de Guyane. Tome 1. Collection du Patrimoine Naturel, Vol. 22. IEBGMNHN, INRA, Min. Env. Paris.

Pompeu, P.D.S., 1999. Diet of pirambeba *Serrasalmus brandtii* Reinhardt (Teleostei, Characidae) in four floodplain lakes in São Francisco river, Brazil. Revista Brasileira de Zoologia, 16, pp.19-26.

Pouilly, M., Lino, F., Bretenoux, J.G. and Rosales, C., 2003. Dietary–morphological relationships in a fish assemblage of the Bolivian Amazonian floodplain. Journal of fish Biology, 62(5), pp.1137-1158.

Pouilly, M., T. Yunoki, C. Rosales, and L. Torres. 2004. Trophic structure of fish assemblages from Mamoré River floodplain lakes (Bolivia). Ecology of Freshwater Fish 13(4): 245-57.

Prudente, B.D.S., Carneiro-Marinho, P., Valente, R.D.M. and Montag, L.F.D.A., 2016. Feeding ecology of *Serrasalmus gouldingi* (Characiformes: Serrasalmidae) in the lower Anapu River region, eastern Amazon, Brazil. Acta Amazonica, 46(3), pp.259-270.

Ramos, I.P., Vidotto-Magnoni, A.P. and Carvalho, E.D., 2008. Influence of cage fish farming on the diet of dominant fish species of a Brazilian reservoir (Tietê River, High Paraná River basin). Acta Limnologica Brasiliensia, 20(3), pp.245-252.

Raposo, R.D.M.G. and Gurgel, H.D.C.B., 2003. Variação da alimentação natural de Serrasalmus spilopleura Kner, 1860 (Pisces, Serrasalmidae) em função do ciclo lunar e das estações do ano na lagoa de Extremoz, Rio Grande do Norte, Brasil. Acta Scientiarum. Animal Sciences, 25(2), pp.267-272.

Röpke, C.P., Ferreira, E. and Zuanon, J., 2014. Seasonal changes in the use of feeding resources by fish in stands of aquatic macrophytes in an Amazonian floodplain, Brazil. Environmental biology of fishes. 97(4): 401-414.

Santos, G. M., S. S. Pinto, and M. Jégu. 1997. Alimentação do pacu-cana, *Mylesinus paraschomburgkii* (Teleostei, Serrasalmidae) em Rios da Amazônia Brasileira. Revista Brasileira de Biologia 57(2): 311-15.

Silva-Camacho, D.D.S., Santos, J.N.D.S., Gomes, R.D.S. and Araújo, F.G., 2014. Ecomorphological relationships among four Characiformes fish species in a tropical reservoir in South-eastern Brazil. Zoologia (Curitiba), 31(1), pp.28-34.

Sório, V.F., Damasceno–Junior, G.A. and Parolin, P., 2014. Dispersal of palm seeds (Bactris glaucescens Drude) by the fish Piaractus mesopotamicus in the Brazilian Pantanal. Ecotropica, 20(1/2), pp.75-82.

Trindade, M.E.D.J. and Jucá-Chagas, R., 2008. Diet of two serrasalmin species, Pygocentrus piraya and Serrasalmus brandtii (Teleostei: Characidae), along a stretch of the rio de Contas, Bahia, Brazil. Neotropical Ichthyology, 6(4), pp.645-650.

Vieira, I. and Géry, J., 1979. Crescimento diferencial e nutrição em *Catoprion mento* (Characoidei). Peixe lepidófago da Amazônia. Acta Amazonica, 9(1), pp.143-146.

Vitorino Júnior, O.B., Agostinho, C.S. and Pelicice, F.M., 2016. Ecology of *Mylesinus paucisquamatus* Jégu & Santos, 1988, an endangered fish species from the rio Tocantins basin. Neotropical Ichthyology, 14(2).

Wantzen, K.M., de Arruda Machado, F., Voss, M., Boriss, H. and Junk, W.J., 2002. Seasonal isotopic shifts in fish of the Pantanal wetland, Brazil. Aquatic Sciences, 64(3), pp.239-251.

Winemiller, K.O., 1989. Ontogenetic diet shifts and resource partitioning among piscivorous fishes in the Venezuelan ilanos. Environmental Biology of fishes, 26(3), pp.177-199.

Zarske, A. and Géry, J., 2008. Revision der neotypischen Gattung *Metynnis* Cope, 1878. II. Beschreibung zweier neuer arten und zum status von *Metynnis goeldii* Eigenmann, 1903 (Teleostei: Characiformes: Serrasalmidae). Vert. Zool, 58, pp.173-196.

Zuluaga-Gómez, M. A., D. B. Fitzgerald, T. Giarrizzo, and K. O. Winemiller. 2016. Morphologic and trophic diversity of fish assemblages in rapids of the Xingu River, a major Amazon tributary and region of endemism. Environmental Biology of Fishes 99(8-9): 647-58.

**S7 - TAXONOMIC RECOMMENDATIONS & MORPHOLOGICAL SYNAPOMORPHIES**

The current study provides robust molecular support for recognizing three major lineages of Serrasalmidae at the subfamilial rank: Colossominae (pacus common to lowland, white water habitats), Myleinae (pacus common to upland clear- and black water habitats), and Serrasalminae (*Metynnis* and piranhas, cosmopolitan). Likewise, recent molecular studies have helped place those taxa in a phylogenetic framework (Freeman et al. 2007; Ortí et al. 2008; Thompson et al. 2014) and uncovered new species-level diversity (Machado et al., 2018). Furthermore, there is strong support for the sister group relationship between Myleinae and Serrasalminae. Those results are consistent with previous phylogenies based on morphological (Cione et al., 2009) and molecular (Ortí et al., 2008; Thompson et al., 2014) data.

Among serrasalmins, the genus *Pristobrycon* Eigenmann 1915 has long been problematic for piranha taxonomy. The genus is considered an artificial (non-monophyletic) assemblage of at least six species divided into two groups, those with a preanal spine (*P. calmoni*, *P. aureus* and *P. eigenmanni*) and those without (*P. careospinus*, *P. maculipinnis* and *P. striolatus*) (Nico et al., 2018). The current study did not include *P. maculipinnis*, but placed the remaining species into three separate groups: ‘*aureus*’ clade (*P. aureus*, *P. careospinus* and *P. eigenmanni*), sister group to monotypic genera *Catoprion* + *Pygopristis* (*P. striolatus*), and the roving *Pristobrycon calmoni*, type species of the genus. Placement of *P. calmoni* varied between the three analyses. The concatenated amino acid analysis supported a sister group relationship between *P. calmoni* and the ‘*maculatus*’ clade, whereas the concatenated nucleotide analysis placed it sister to the ‘*maculatus*’ + ‘*rhombeus*’ clade. The MSC analysis placed P*. calmoni* sister to the ‘*rhombeus*’ clade. The simplest way to resolve the status of *Pristobrycon* is to expand the genus *Serrasalmus* to include *P.* *calmoni* as well as *P. aureus*, *P. careospinus*, *P. eigenmanni*. That said, *Pristobrycon striolatus* and the cryptic *P. scapularis* (Andrade et al., 2019) are not closely related to *Serrasalmus* and warrant a new generic name.

Among myleins, the genus *Tometes* is newly problematic. The genus includes seven species distributed in rivers draining the Guiana Shield into the Orinoco and Negro (*T. makue*), Amazon (*T. camunani* and *T. trilobatus* in part), and coastal rivers from the Maroni to the Araguari (*T. lebaili* and *T. trilobatus* in part) as well as rivers draining the Brazilian Shield into the Amazon (*T. ancylorhynchus*, *T.* *kranponha* and *T. siderocarajensis*) (Andrade et al. 2017). In our analysis, two species from rivers draining the Brazilian Shield, *T. ancylorhynchus* and *T.* *kranponha*, are more closely related to species of *Mylesinus* and *Myleus* than to *Tometes* from coastal rivers draining the Guiana Shield, *T. lebaili* and *T. trilobatus* (type species). An analysis of an extensive dataset of DNA barcodes also failed support the monophyly of *Tometes* (Machado et al. 2018). The polyphyly of *Tometes* warrants further testing.

Finally, our analyses fail to support the exclusive monophyly of *Myloplus,* a result consistent with other molecular studies (Ortí et al. 2008; Thompson et al. 2014; Machado et al. 2018). Species currently assigned to *Myloplus* form up to six different lineages within Myleinae. For examples, *Myloplus schomburgkii* is the sister taxon to the monotypic *Ossubtus*, and *M.* cf. *lucienae* is the sister taxon to Brazilian Shield *Tometes*. *Myloplus arnoldi* is the sister taxon to a clade composed of *M. lucienae*, *M. planquettei*, Brazilian Shield *Tometes*, and species of *Mylesinus* and *Myleus*. *Myloplus planquettei* nests within *Myleus* and members of this clade share number of morphological characteristics, notably an elongate cranium and wide-cusped teeth. *Myloplus rhomboidalis* is the sister taxon to all other myleins except *Acnodon*. *Myloplus rhomboidalis* (Cuvier 1818) is the type species of nominal subgenus *Prosomyleus* Géry 1972. Therefore, we recommend resurrecting *Prosomyleus* from the synonymy of *Myloplus* and elevating it to generic rank for species *P. rhomboidalis*.

The clade of *Myloplus* inclusive of the type species *M. asterias* (Müller & Troschel 1844) groups congeners *M.* cf. *lobatus*, *M. rubripinnis*, *M. taphorni*, *M. ternetzi*, and *M.* cf. *torquatus*. All of our analyses also nested *Utiaritichthys* within this clade of true *Myloplus*. An analysis of DNA barcodes similarly nested *Utiaritichthys* well within *Myloplus* (Machado et al., 2018). *Utiaritichthys* is distinguished in part by its elongate bauplan, a feature it shares with *Myloplus asterias*. Therefore, we recommend the synonymization of *Utiaritichthys* Miranda Ribeiro 1937 with *Myloplus* Gill 1896.

**Family Serrasalmidae: Günther, 1864**

*Morphological synapomorphies*. The monophyly of this family is based on morphological synapomorphies proposed by Machado-Allison (1983, 1985) and by Buckup (1998). More recently, Kolmann et al. (2018) proposed the presence of a serrate, mid-ventral keel as being a synapomorphy for Serrasalmidae and subsequently, Kolmann et al. (2019) also proposed that unilateral tooth replacement is also a synapomorphy for serrasalmids.

*Comment*. The subfamilies recognized in Serrasalmidae, Serrasalminae and Myleinae, follow Buckup (1998b) and Machado-Allison (1982, 1983, 1985). Myleinae and Serrasalminae were first proposed by Eigenmann (1915), with the former diagnosed by having two rows of premaxillary teeth and the latter having a single row of premaxillary teeth. Machado-Allison (1983) later modified these definitions, by proposing that the Myleinae be further distinguished by often having one pair of symphyseal teeth on the dentary and the Serrasalminae having tricuspid teeth. The major difference between Eigenmann’s definition of the subfamilies *vs*. Machado-Allison’s is that the former included *Catoprion* and *Metynnis* within the Serrasalminae.

*Classification of Serrasalmidae*

A new classification of suprageneric groups within Serrasalmidae is proposed based on the current molecular analysis (Fig. 2).

**Family** Serrasalmidae Bleeker 1859

**Subfamily** Colossominae new subfamily*

**Included valid nominal genera:** *Colossoma* Eigenmann & Kennedy 1903, *Mylossoma* Eigenmann & Kennedy 1903, and *Piaractus* Eigenmann 1903

**Subfamily** Myleinae Eigenmann 1903

**Included valid nominal genera:** *Acnodon* Eigenmann 1903, *Mylesinus* Valenciennes 1850, *Myleus* Müller & Troschel 1844, *Myloplus* Gill 1896 (includes *Utiaritichthys* Miranda Ribeiro 1937), *Ossubtus* Jégu 1992, and *Tometes* Valenciennes 1850

**Subfamily** Serrasalminae Bleeker 1859

**Included valid nominal genera:** *Catoprion* Müller & Troschel 1844, *Metynnis* Cope 1878, *Prosomyleus* Géry 1972, Pygocentrus, *Pygopristis* Müller & Troschel 1844, and *Serrasalmus* Lacepède 1803 (includes *Pristobrycon* Eigenmann 1915).

*Putative synapomorphies uniting *Mylossoma, Colossoma,* & *Piaractus*:

- generally with > 40 abdominal serrae (Machado-Allison, 1983; Kolmann et al., 2018)
- The dorsal fin not preceded by a spinous process continuous with the first pterygiophore (Machado-Allison, 1983)
- The intercalar bone is large and firmly attached to the neurocranium (Machado-Allison, 1983)
- Absence of a humeral hiatus in the anterolateral muscular body wall (Machado-Allison, 1983)
- Well-developed and elongate pterotic spine (Machado-Allison, 1983)
- Robust frontal bones with well-developed lateral extensions, which form a deep dilatator fossa (Machado-Allison, 1983)
- Robust, laterally-expanded mesethmoid (Machado-Allison, 1983)
