## Supplementary material for "Phylogenomics of piranhas and pacus (Serrasalmidae) uncovers how convergent diets obfuscate traditional morphological taxonomy": S8

1 Table S8. List of taxa, voucher specimens, and DNA sequences analyzed. Museum codes follow Sabaj (2018).

| | Superfamily / Family / Terminal taxon | In Tree As: | Voucher musem and catalog number | Voucher tag or other identifer | Tissue catalog number | Country | Basin: Sub-basins |
|---|---|---|---|---|---|---|---|
| 2 | Superfamily | | | | | | |
| 3 | **Family** | | | | | | |
| 4 | Terminal taxon | **In Tree As:** | **Voucher musem and catalog number** | **Voucher tag or other identifer** | **Tissue catalog number** | **Country** | **Basin: Sub-basins** |
| 5 | Alestoidea Cockerell 1910 | | | | | | |
| 6 | **Alestidae Cockerell 1910** | | | | | | |
| 7 | *Alestopetersius hilgendorfi* (Boulenger 1899) | | **AMNH 242460** | **AMNH 034-3321** | | **Democratic Republic of the Congo** | **Lac Ikenge** |
| 8 | *Alestopetersius tumbensis* (Hoedeman 1951) | | **AMNH 247033** | **AMNH 052-5189** | | **Democratic Republic of the Congo** | **Congo River** |
| 9 | *Bathyaethiops greeni (* Fowler 1949) | | **AMNH 242750** | **AMNH 024-2395** | | **Democratic Republic of the Congo** | **Luilaka River** |
| 10 | *Brycinus grandisquamis* (Boulenger 1899) | | **AMNH 257862** | **AMNH 115-11429** | | **Democratic Republic of the Congo** | **Ndolo** |
| 11 | *Bryconalestes longipinnis* (Günther 1864) | | **AUFT 59705** | **Guinea-0560** | **AUFT 5548** | **Republic of Guinea** | **Nianda River : St.Paul River** |
| 12 | *Hydrocynus goliath* Boulenger 1898 | | **AMNH 239463** | **AMNH 022-2138** | | **Republic of the Congo** | **Congo, near Brazzaville** |
| 13 | *Micralestes acutidens* (Peters 1852) | | **AUFT 51513** | **AFR0013** | **AUFT 5674** | **Democratic Republic of the Congo** | **Congo River : Atlantic Ocean** |
| 14 | *Rhabdalestes septentrionalis* (Boulenger 1911) | | **AMNH 257107** | **AMNH 113-11282** | | **Guinea** | **Mong River** |
| 15 | **Hepsetidae Hubbs 1939** | | | | | | |
| 16 | *Hepsetus odoe* (Bloch 1794) | | **AUFT 56869** | **JWA192** | **AUFT 5790** | **Cameroon** | **Congo River** |
| 17 | Erythrinoidea Valenciennes 1847 | | | | | | |
| 18 | **Erythrinidae Valenciennes 1847** | | | | | | |
| 19 | *Hoplerythrinus unitaeniatus* (Spix & Agassiz 1829) | | **AUFT 48549** | **G07-341** | **AUFT 4875** | **Guyana** | **Rio Branco : Rio Negro : Rio Amazonas** |
| 20 | *Hoplias aimara* (Valenciennes 1847) | | **AUFT 47950** | **G07-207** | **AUFT 4938** | **Guyana** | **Essequibo River** |
| 21 | Characoidea Latreille 1825 | | | | | | |
| 22 | **Chalceidae Fowler 1958** | | | | | | |
| 23 | *Chalceus epakros* Zanata & Toledo-Piza 2004 | | **AUFT 48194** | **G07-711** | **AUFT 3601** | **Guyana** | **Rupununi River : Essequibo River** |
| 24 | **Characidae Latreille 1825** | | | | | | |
| 25 | *Nematobrycon palmeri* Eigenmann 1911 | | tba ROM | **MK-057** | | **Colombia** | **Amazonas** |
| 26 | *Petitella georgiae* Géry & Boutière 1964 | | tba ROM | **MK-060** | | **Peru** | **Amazonas: Nanay** |
| 27 | Curimatoidea Gill 1858 | | | | | | |
| 28 | **Anostomidae Günther 1864** | | | | | | |

| | | | | | | |
|---|---|---|---|---|---|---|
| 29 | *Abramites hypselonotus* (Günther 1868) | **G. Ortí** | **GO-77** | **GO-77** | **locality unknown** | **locality unknown** |
| 30 | *Anostomus ternetzi* Fernández-Yépez 1949 | **AUFT 48209** | **G07-749** | **AUFT 3075** | **Guyana** | **Rupununi River : Essequibo River** |
| 31 | *Laemolyta proxima* (Garman 1890) | **AUFT 47859** | **G07-102** | **AUFT 3059** | **Guyana** | **Rupununi River, at Kwatamang landing** |
| 32 | *Leporinus brunneus* Myers 1950 | **AUFT 43799** | **V5389** | **AUFT 3071** | **Venezuela** | **Río Negro : Amazon River** |
| 33 | *Leporinus ortomaculatus* Garavello 2000 | **AUFT 43964** | **V5582** | **AUFT 3065** | **Venezuela** | **Río Orinoco** |
| 34 | *Leporinus striatus* Kner 1858 | **AUFT 43977** | **V5587** | **AUFT 3064** | **Venezuela** | **Río Orinoco** |
| 35 | *Petulanos spiloclistron* (Winterbottom 1974) | **AUFT 44717** | **G5207** | **AUFT 3057** | **Guyana** | **Rio Branco : Rio Negro** |
| 36 | *Pseudanos winterbottomi* Sidlauskas & Santos 2005 | **AUFT 51124** | **V5288** | **AUFT 3084** | **Venezuela** | **Río Ventuari : Río Orinoco** |
| 37 | *Schizodon vittatus* (Valenciennes 1850) | **AUM 35502** | **T2417** | | **Guyana** | **Essequibo: Rupununi** |
| 38 | *Synaptolaemus latofasciatus* (Steindachner 1910) | **AUFT 43977** | **V5587** | **AUFT 3064** | **Venezuela** | **Río Orinoco** |
| 39 | **Chilodontidae Eigenmann 1903** | | | | | |
| 40 | *Caenotropus labyrinthicus* (Kner 1858) | **AUFT 43784** | **V5381** | **AUFT 3096** | **Venezuela** | **Río Negro : Amazon River** |
| 41 | **Curimatidae Gill 1858** | | | | | |
| 42 | *Curimata roseni* Vari 1989 | **ANSP 189094** | **V071** | **ANSP t1311** | **Venezuela** | **Orinoco: Ventuari** |
| 43 | *Curimatopsis cryptica* Vari 1982 | **ANSP 189091** | **7057** | **ANSP t1308** | **Suriname** | **Suriname: Coropinae** |
| 44 | *Cyphocharax abramoides* (Kner 1858) | **AUM 42957** | **V5411** | | **Venezuela** | **Orinoco** |
| 45 | *Psectrogaster ciliata* (Müller & Troschel 1844) | **ANSP 189093** | **V072** | **ANSP t1309** | **Venezuela** | **Orinoco: Ventuari** |
| 46 | **Cynodontidae Eigenmann 1903** | | | | | |
| 47 | *Cynodon gibbus* (Spix & Agassiz 1829) | **ANSP 182523** | **P6002** | **ANSP t4570** | **Peru** | **Amazonas: Nanay** |
| 48 | *Cynodon gibbus* (Spix & Agassiz 1829) | **ROM (no voucher)** | **ROM T11775** | | **Venezuela** | **Rio Cinaruco** |
| 49 | *Cynodon gibbus* (Spix & Agassiz 1829) | **AUFT 44336** | **G5013** | **AUFT 3244** | **Guyana** | **Essequibo River** |
| 50 | *Cynodon meionactis* Géry, Le Bail & Keith 1999 | **ANSP 189129** | **7029** | **ANSP t1597** | **Suriname** | **Maroni: Litanie** |
| 51 | *Cynodon meionactis* Géry, Le Bail & Keith 1999 | **ROM (no voucher)** | **ROM T19037** | | **Suriname** | **Corantijn River** |
| 52 | *Cynodon septenarius* Toledo-Piza 2000 | **ROM 101254** | **ROM T21832** | | **Guyana** | **Essequibo: Rewa River** |
| 53 | *Cynodon septenarius* Toledo-Piza 2000 | **UMMZ 250675** | **UMMZ T00852** | | **Guyana** | **Essequibo: Rewa River** |
| 54 | *Hydrolycus armatus* (Jardine 1841) | **ANSP 191148** | **BO6116** | **ANSP t3931** | **Venezuela** | **Orinoco: Ventuari** |
| 55 | *Hydrolycus armatus* (Jardine 1841) | **ROM 088356** | **ROM T07147** | | **Peru** | **Amazonas** |

| | | | | | | | |
|---|---|---|---|---|---|---|---|
| 56 | *Hydrolycus armatus* (Jardine 1841) | | **ROM 101263** | **ROM T21613** | | **Guyana** | **Essequibo: Mazaruni River** |
| 57 | *Hydrolycus scomberoides* (Cuvier 1819) | | **ROM 086143** | **ROM T06905** | | **Guyana** | **Essequibo: Rupununi** |
| 58 | *Hydrolycus cf scomberoides* (Cuvier 1819) | | **ROM 088358** | **ROM T10758** | | **Peru** | **Amazonas** |
| 59 | *Hydrolycus tatauaia* Toledo-Piza, Menezes & Santos 1999 | | **ANSP 182629** | **P6337** | **ANSP t1804** | **Peru** | **Amazonas: Nanay** |
| 60 | *Hydrolycus wallacei* Toledo-Piza, Menezes & Santos 1999 | | **ANSP 182425** | **P4316** | **ANSP t1600** | **Venezuela** | **Orinoco** |
| 61 | *Hydrolycus* cf. *wallacei* Toledo-Piza, Menezes & Santos 1999 | | **PUJ** | | | **Colombia** | **Orinoco: Vaupes** |
| 62 | *Rhaphiodon vulpinus* Spix & Agassiz 1829 | | **ANSP 198993** | **t4845** | **ANSP t13122** | **Venezuela** | **Orinoco: Apure** |
| 63 | **Hemiodontidae Bleeker 1859** | | | | | | |
| 64 | *Anodus elongatus* Agassiz 1829 | | **ANSP 182300** | **P6248** | **ANSP t4542** | **Peru** | **Amazonas** |
| 65 | *Argonectes longiceps* (Kner 1858) | | **ANSP 191353** | **BO6102** | **ANSP t4440** | **Venezuela** | **Orinoco: Ventuari** |
| 66 | *Argonectes longiceps* (Kner 1858) | | **ROM 093581** | **ROM T09783** | | **Venezuela** | **Caño Tigre** |
| 67 | *Bivibranchia bimaculata* Vari 1985 | | **ANSP 189149** | **6870** | **ANSP t1823** | **Suriname** | **Maroni: Lawa** |
| 68 | *Bivibranchia fowleri* (Steindachner 1908) | | **LIA 5666** | **t5701** | **ANSP t14127** | **Brazil** | **Amazonas: Xingu** |
| 69 | *Hemiodus* cf. *gracilis* Günther 1864 | | tba ROM | | | **Colombia** | **Amazonas** |
| 70 | *Hemiodus quadrimaculatus* Pellegrin 1909 | | **ROM 096949** | **ROM T18007** | | **Guyana** | **Konawaruk River** |
| 71 | *Hemiodus unimaculatus* (Bloch 1794) | | **ANSP 195843** | **t2436** | **ANSP t10589** | **Brazil** | **Amazonas: Xingu** |
| 72 | *Micromischodus sugillatus* Roberts 1971 | | **FMNH 189039** | **UCFJDBG 14101353** | | **Brazil** | **Arapiuns River** |
| 73 | **Parodontidae Eigenmann 1910** | | | | | | |
| 74 | *Parodon guyanensis* Géry 1960 | | **AUFT 48207** | **G07-709** | **AUFT 3605** | **Guyana** | **Rupununi River : Essequibo River** |
| 75 | *Parodon orinocensis* (Bonilla, Machado-Allison, Silvera, Chernoff, López & Lasso 1999) | | **AUFT 43666** | **V5251** | **AUFT 3957** | **Venezuela** | **Río Negro : Río Amazonas** |
| 76 | **Prochilodontidae Eigenmann 1909** | | | | | | |
| 77 | *Ichthyoelephas longirostris* (Steindachner 1879) | | **ANSP 192865** | **6609** | **ANSP t1305** | **Colombia** | **Magdalena** |
| 78 | *Prochilodus scrofa* Steindachner 1881 | | **AUFT 46587** | **P6077** | **AUFT 3735** | **Peru** | **Río Marañón** |
| 79 | *Semaprochilodus varii* Castro 1988 | | **ANSP 187435** | **6929** | ANSP t1240 | **Suriname** | **Maroni (Marowijne): Lawa** |
| 80 | **Serrasalmidae Bleeker 1859** | **In Figure 2 as:** | | | | | |
| 81 | *Acnodon normani* Gosline 1951 | ***Acnodon normani 1*** | **ANSP 199545** | **B1548** | **ANSP t5742** | **Brazil** | **Amazonas: Xingu** |

| | | | | | | | |
|---|---|---|---|---|---|---|---|
| 82 | *Acnodon normani* Gosline 1951 | ***Acnodon normani 2*** | tba ROM | **T26642** | | **Brazil** | **Amazonas: Tocantins** |
| 85 | *Acnodon oligacanthus* (Müller & Troschel 1844) | ***Acnodon oligacanthus 1*** | **MNHN 1998-1585** | | | **Suriname** | **Maroni (Marowijne)** |
| 83 | *Acnodon oligacanthus* (Müller & Troschel 1844) | ***Acnodon oligacanthus 2*** | **ROM 100851** | **t19957** | | **Suriname** | **Maroni (Marowijne)** |
| 84 | *Acnodon oligacanthus* (Müller & Troschel 1844) | ***Acnodon oligacanthus 3*** | **ROM 100851** | **t19959** | | **Suriname** | **Maroni (Marowijne)** |
| 86 | *Catoprion mento* (Cuvier 1819) | ***Catoprion mento 1*** | **ROM 85951** | **t06600** | | **Guyana** | **Pirara** |
| 87 | *Catoprion mento* (Cuvier 1819) | ***Catoprion mento 2*** | **ROM 95239** | **t17406** | | **Brazil** | **Amazonas: Tapajós** |
| 88 | *Catoprion mento* (Cuvier 1819) | ***Catoprion mento 3*** | **UMMZ** | **t01162** | | **Guyana** | |
| 89 | *Colossoma macropomum* (Cuvier 1816) | ***Colossoma macropomum 1*** | **ANSP 198904** | **t4874** | **ANSP t12983** | **Venezuela** | **Orinoco: Apure** |
| 90 | *Colossoma macropomum* (Cuvier 1816) | ***Colossoma macropomum 2*** | **ANSP 198904** | **t4876** | **ANSP t12984** | **Venezuela** | **Orinoco: Apure** |
| 93 | *Colossoma macropomum* (Cuvier 1816) | ***Colossoma macropomum 3*** | **INPA 10149** | **GO-216** | | **Brazil** | **Rio Solimoes** |
| 91 | *Colossoma macropomum* (Cuvier 1816) | ***Colossoma macropomum 4*** | **PUJ tbd** | **MK-012** | | **Colombia** | **Amazonas** |
| 92 | *Colossoma macropomum* (Cuvier 1816) | ***Colossoma macropomum 5*** | **PUJ tbd** | | | **Colombia** | **Amazonas** |
| 94 | *Metynnis altidorsalis* Ahl 1923 | ***Metynnis altidorsalis 1*** | **ANSP 198246** | **t3646** | **ANSP t12536** | **Brazil** | **Amazonas: Xingu** |
| 95 | *Metynnis* cf. *argenteus* Ahl 1923 | ***Metynnis cf argenteus 1*** | **ROM 86134** | **t06595** | | **Suriname** | **Corantijn** |
| 96 | *Metynnis fasciatus* Ahl 1931 | ***Metynnis fasciatus 1*** | tba ROM | **MK-18-012** | | **Peru** | **Amazonas: Nanay** |
| 97 | *Metynnis fasciatus* Ahl 1931 | ***Metynnis fasciatus 2*** | tba ROM | **MK-18-014** | | **Peru** | **Amazonas: Nanay** |
| 98 | *Metynnis* cf. *guaporensis* Eigenmann 1915 | ***Metynnis cf guaporensis*** | **UMMZ 250562** | **UMMZ T00683** | | **Guyana** | **Essequibo: Rewa River** |
| 100 | *Metynnis hypsauchen* (Müller & Troschel 1844) | ***Metynnis hypsauchen 1*** | **ROM 87023** | **t8260** | | **Guyana** | **Berbice** |
| 99 | *Metynnis hypsauchen* (Müller & Troschel 1844) | ***Metynnis hypsauchen 2*** | **ANSP 199545** | **t1278** | **ANSP t8923** | **Brazil** | **Amazonas: Xingu** |
| 103 | *Metynnis lippincottianus* (Cope 1870) | ***Metynnis lippicottiaus 1*** | tba ROM | **MK-18-B** | | **Peru** | **Amazonas: Nanay** |
| 101 | *Metynnis lippincottianus* (Cope 1870) | ***Metynnis lippicottiaus 2*** | **INPA 47353** | **t3853** | **ANSP t12699** | **Brazil** | **Amazonas: Xingu** |
| 102 | *Metynnis lippincottianus* (Cope 1870) | ***Metynnis lippicottiaus 3*** | **INPA 47353** | **t3854** | **ANSP t12700** | **Brazil** | **Amazonas: Xingu** |
| 104 | *Metynnis* cf. *longipinnis* Zarske & Géry 2008 | ***Metynnis cf longipinnis*** | **UMMZ 250564** | **UMMZ T00674** | | **Guyana** | **Essequibo: Rewa River** |
| 105 | *Metynnis luna* Cope 1878 | ***Metynnis guaporensis 1*** | **ANSP 198808** | **t3283** | **ANSP t12210** | **Brazil** | **Amazonas: Xingu** |
| 106 | *Metynnis luna* or *guaporensis* Cope 1878 | ***Metynnis guaporensis 2*** | **INPA 47413** | **t3281** | **ANSP t12211** | **Brazil** | **Amazonas: Xingu** |
| 107 | *Metynnis* cf. *luna* Cope 1878 | ***Metynnis cf luna 1*** | **ROM 86118** | **t06561** | | **Guyana** | **Essequibo** |
| 108 | *Metynnis* cf. *luna* Cope 1878 | ***Metynnis cf luna 2*** | **ROM 86118** | **t06560** | | **Guyana** | **Essequibo** |

| | | | | | | | |
|---|---|---|---|---|---|---|---|
| 109 | *Metynnis maculatus* (Kner 1858) | ***Metynnis maculatus 1*** | **UMMZ 250564** | **UMMZ T00673** | | **Guyana** | **Essequibo: Rewa River** |
| 110 | *Metynnis maculatus* (Kner 1858) | ***Metynnis maculatus 2*** | **UMMZ 250564** | **UMMZ T00675** | | **Guyana** | **Essequibo: Rewa River** |
| 111 | *Metynnis* sp. | ***Metynnis sp.*** | **UMMZ 252270** | **UMMZ T00672** | | **Guyana** | **Essequibo: Rewa River** |
| 112 | *Mylesinus paucisquamatus* Jégu & Santos 1988 | ***Mylesinus paucisquamatus 1*** | tba ROM | **MK-18-003** | | **Brazil** | **Amazonas: Jari** |
| 113 | *Mylesinus paucisquamatus* Jégu & Santos 1988 | ***Mylesinus paucisquamatus 2*** | tba ROM | **MK-18-004** | | **Brazil** | **Amazonas: Jari** |
| 114 | *Myleus* cf. *knerii* (Steindachner 1881) | ***Myleus cf. knerii*** | **ROM 86416** | **t06997** | | **Guyana** | **Essequibo: Rupununi** |
| 115 | *Myleus pacu* (Jardine 1841) | ***Myleus pacu 1*** | **AUFT 47855** | **G07-093** | **AUFT 3042** | **Guyana** | **Essequibo River** |
| 119 | *Myleus* cf. *setiger* Müller & Troschel 1844 | ***Myleus cf setiger 1*** | **PUJ** | | | **Venezuela** | **Parucito:Ventuari** |
| 120 | *Myleus* cf. *setiger* Müller & Troschel 1844 | ***Myleus cf setiger 2*** | **UMMZ 250650** | **UMMZ T00829** | | **Guyana** | **Essequibo: Rewa River** |
| 116 | *Myleus setiger* Müller & Troschel 1844 | ***Myleus setiger 1*** | **ANSP 197912** | **t3546** | **t12474** | **Brazil** | **Amazonas: Xingu** |
| 118 | *Myleus setiger* Müller & Troschel 1844 | ***Myleus setiger 2*** | **ROM** | **t9693** | | **Guyana** | **Essequibo: Rupununi** |
| 117 | *Myleus setiger* Müller & Troschel 1844 | ***Myleus setiger 3*** | **ROM** | **t6914** | | **Guyana** | **Essequibo: Rupununi** |
| 121 | *Myloplus rubripinnis* (Müller & Troschel 1844) | ***Myloplus arnoldi 1*** | **ANSP 195980** | **t2771** | **t10913** | **Brazil** | **Amazonas: Xingu: Iriri** |
| 122 | *Myloplus arnoldi* Ahl 1936 | ***Myloplus arnoldi 2*** | **ANSP 193058** | **B2132** | **t5915** | **Brazil** | **Amazonas: Xingu** |
| 123 | *Myloplus asterias* (Müller & Troschel 1844) | ***Myloplus asterias 1*** | **ROM 97246** | **t18308** | | **Guyana** | **Berbice** |
| 124 | *Myloplus asterias* (Müller & Troschel 1844) | ***Myloplus asterias 2*** | **ROM 101260** | **t21670M** | | **Guyana** | **Essequibo: Mazaruni** |
| 125 | *Myloplus* cf. *lobatus* (Valenciennes 1850) | ***Myloplus cf lobatus 1*** | tba ROM | **ROM t26644** | **MK-039** | **Peru** | **Amazonas: Nanay** |
| 126 | *Myloplus lucienae* Andrade, Ota, Bastos & Jégu 2016 | ***Myloplus cf lucienae*** | | **AUM p4661** | | **locality unknown** | **locality unknown** |
| 127 | *Myloplus planquettei* Jégu, Keith & Le Bail 2003 | ***Myloplus planquettei 1*** | **ROM 102376** | **t23037** | | **Suriname** | **Saramacca** |
| 128 | *Myloplus planquettei* Jégu, Keith & Le Bail 2003 | ***Myloplus planquettei 2*** | **ROM 102441** | **t23116** | | **Suriname** | **Saramacca** |
| 133 | *Myloplus* cf. *rhomboidalis* (Cuvier 1818) | ***Myloplus cf rhomboidalis*** | **PUJ tbd** | **ROM t26513** | **MK-010** | **Colombia** | **Orinoco: Vaupes** |
| 131 | *Myloplus rhomboidalis* (Cuvier 1818) | ***Myloplus rhomboidalis 1*** | **ROM 101260** | **t21669F** | | **Guyana** | **Essequibo: Mazaruni** |
| 132 | *Myloplus rhomboidalis* (Cuvier 1818) | ***Myloplus rhomboidalis 2*** | **ROM 102444** | **t23210** | | **Suriname** | **Saramacca** |
| 129 | *Myloplus rhomboidalis* (Cuvier 1818) | ***Myloplus rhomboidalis 3*** | **ROM 98019** | **t18919** | | **Suriname** | **Corantijn** |
| 130 | *Myloplus rhomboidalis* (Cuvier 1818) | ***Myloplus rhomboidalis 4*** | **ROM 100574** | **t20182** | | **Suriname** | **Corantijn** |
| 134 | *Myloplus rubripinnis* (Müller & Troschel 1844) | ***Myloplus rubripinnis 1*** | **AUM 43899** | **V5553** | **AUFT 3051** | **Venezuela** | **Orinoco** |
| 135 | *Myloplus rubripinnis* (Müller & Troschel 1844) | ***Myloplus rubripinnis 2*** | **ROM 96256** | **t16811** | | **Guyana** | **Essequibo: Kuyuwini** |

| | | | | | | | |
|---|---|---|---|---|---|---|---|
| 136 | *Myloplus rubripinnis* (Müller & Troschel 1844) | ***Myloplus rubripinnis 3*** | **ROM 96256** | **t16830** | | **Guyana** | **Essequibo: Kuyuwini** |
| 139 | *Myloplus schomburgkii* (Jardine 1841) | ***Myloplus schmoburgkii 1*** | **ROM 88293** | **t9434** | | **Venezuela** | **Orinoco: Atabapo** |
| 137 | *Myloplus schomburgkii* (Jardine 1841) | ***Myloplus schmoburgkii 2*** | **ANSP 194779** | **t0546** | **t8191** | **Brazil** | **Amazonas: Xingu** |
| 138 | *Myloplus schomburgkii* (Jardine 1841) | ***Myloplus schmoburgkii 3*** | **ROM 88932** | **t7186** | | **Peru** | **Amazonas: Nanay** |
| 141 | *Myloplus taphorni* Andrade, López-Fernández & Liverpool 2019 | ***Myloplus taphorni 1*** | **ROM 101261** | **t21742** | | **Guyana** | **Essequibo: Mazaruni** |
| 140 | *Myloplus taphorni* Andrade, López-Fernández & Liverpool 2019 | ***Myloplus taphorni 2*** | **ROM 101261** | **t21730** | | **Guyana** | **Essequibo: Mazaruni** |
| 142 | *Myloplus taphorni* Andrade, López-Fernández & Liverpool 2019 | ***Myloplus taphorni 3*** | **ROM 102709** | **t21809** | | **Guyana** | **Essequibo: Mazaruni** |
| 143 | *Myloplus ternetzi* (Norman 1929) | ***Myloplus ternetzi 1*** | **ANSP 188689** | **6984** | **t1626** | **Suriname** | **Maroni: Lawa** |
| 144 | *Myloplus ternetzi* (Norman 1929) | ***Myloplus ternetzi 2*** | **ROM 97890** | **t18664** | | **Suriname** | **Maroni** |
| 145 | *Myloplus* cf. *torquatus* (Kner 1858) | ***Myloplus cf torquatus*** | **PUJ tbd** | **ROM t26429** | **MK-22** | **Colombia** | **Orinoco: Vaupes** |
| 146 | *Mylossoma* sp. | ***Mylossoma sp. 1*** | **AUM 50347** | **P6074** | **AUFT 3733** | **Peru** | **Amazonas: Marañón** |
| 147 | *Mylossoma albiscopum* (Cope 1872) | ***Mylossoma albiscopum 1*** | **PUJ tbd** | | | **Colombia** | **Amazonas** |
| 149 | *Mylossoma albiscopum* (Cope 1872) | ***Mylossoma duriventre 1*** | **ANSP 198905** | **t4877** | **t12979** | **Venezuela** | **Orinoco: Apure** |
| 148 | *Mylossoma aureum* (Spix & Agassiz 1829) | ***Mylossoma aureum 1*** | **ANSP 178147** | **1725** | **t3861** | **Peru** | **Amazonas: Napo** |
| 152 | *Ossubtus xinguense* Jégu 1992 | ***Ossubtus xinguense 1*** | **ANSP 194758** | **t0503** | **t8148** | **Brazil** | **Amazonas: Xingu** |
| 150 | *Ossubtus xinguense* Jégu 1992 | ***Ossubtus xinguense 2*** | **INPA 13195** | **GO-253** | | **Brazil** | **Amazonas: Xingu: Iriri** |
| 151 | *Ossubtus xinguense* Jégu 1992 | ***Ossubtus xinguense 3*** | **INPA 38073** | **B2147** | **t5930** | **Brazil** | **Amazonas: Xingu** |
| 153 | *Piaractus brachypomus* (Cuvier 1818) | ***Piaractus brachypomus 1*** | **PUJ** | **ROM t26507** | | **Colombia** | **Amazonas** |
| 154 | *Piaractus brachypomus* (Cuvier 1818) | ***Piaractus brachypomus 2*** | **ROM 79182** | **ROM t00349** | | **locality unknown** | **locality unknown** |
| 155 | *Piaractus mesopotamicus* (Holmberg 1887) | ***Piaractus mesopotamicus 1*** | **ANSP 182451** | **A5113** | **ANSP t511** | **Argentina** | **Paraná** |
| 156 | *Pristobrycon aureus* (Spix & Agassiz 1829) | ***Pristobrycon aureus 1*** | **UMMZ 250945** | **UMMZ T01191** | | **Guyana** | **Essequibo: Rupununi** |
| 158 | *Pristobrycon aureus* (Spix & Agassiz 1829) | ***Pristobrycon aureus 2*** | **UMMZ 251259** | **UMMZ T01232** | | **Guyana** | **Essequibo: Rupununi** |
| 157 | *Pristobrycon aureus* (Spix & Agassiz 1829) | ***Pristobrycon aureus 3*** | **UMMZ 251259** | **UMMZ T01231** | | **Guyana** | **Essequibo: Rupununi** |
| 159 | *Pristobrycon calmoni* (Steindachner 1908) | ***Pristobrycon calmoni 1*** | **ROM 67437** | **t05407** | | **Guyana** | **Waini River** |
| 160 | *Pristobrycon careospinus* Fink & Machado-Allison 1992 | ***Pristobrycon cf careospinus 1*** | **PUJ** | **MK-002** | | **Colombia** | **Orinoco: Vaupes** |
| 166 | *Pristobrycon striolatus* (Steindachner 1908) | ***Pristobrycon striolatus 1*** | **MNHN 1988-1573** | | | **Suriname** | **Maroni** |
| 165 | *Pristobrycon striolatus* (Steindachner 1908) | ***Pristobrycon striolatus 2*** | **ANSP 188672** | **7031** | **t1631** | **Suriname** | **Maroni: Litanie** |

| | | | | | | | |
|---|---|---|---|---|---|---|---|
| 163 | *Pristobrycon striolatus* (Steindachner 1908) | ***Pristobrycon striolatus 3*** | **ANSP 197515** | **t2333** | **t10494** | **Brazil** | **Amazonas: Xingu** |
| 164 | *Pristobrycon striolatus* (Steindachner 1908) | ***Pristobrycon striolatus 4*** | **ANSP 198809** | **t3261** | **t12228** | **Brazil** | **Amazonas: Xingu** |
| 167 | *Pygocentrus cariba* (Humboldt 1821) | ***Pygocentrus cariba 1*** | **ANSP 198959** | **t4716** | **ANSP t13040** | **Venezuela** | **Orinoco: Apure** |
| 168 | *Pygocentrus cariba* (Humboldt 1821) | ***Pygocentrus cariba 2*** | **ANSP 198959** | **t4717** | **ANSP t13041** | **Venezuela** | **Orinoco: Apure** |
| 169 | *Pygocentrus nattereri* Kner 1858 | ***Pygocentrus nattereri 1*** | **ANSP 197517** | **t2340** | **ANSP t10501** | **Brazil** | **Amazonas: Xingu** |
| 170 | *Pygocentrus nattereri* Kner 1858 | ***Pygocentrus nattereri 2*** | **ROM 85941** | **t06527** | | **Guyana** | **Amazonas: Negro** |
| 171 | *Pygocentrus piraya* (Cuvier 1819) | ***Pygocentrus piraya 1*** | tba ROM | | | **Brazil** | **São Francisco** |
| 172 | *Pygocentrus piraya* (Cuvier 1819) | ***Pygocentrus piraya 2*** | tba ROM | | | **Brazil** | **São Francisco** |
| 173 | *Pygopristis denticulata* (Cuvier 1819) | ***Pygopristis denticulata 1*** | **ANSP 206125** | **T09865** | | **Venezuela** | **Orinoco** |
| 174 | *Pygopristis denticulata* (Cuvier 1819) | ***Pygopristis denticulata 2*** | **G. Ortí** | **GO-412** | | **locality unknown** | **locality unknown** |
| 175 | *Pygopristis denticulata* (Cuvier 1819) | ***Pygopristis denticulata 3*** | **UMMZ 250939** | **UMMZ T01161** | | **Guyana** | **Essequibo: Rupununi** |
| 176 | *Serrasalmus altispinis* Merckx, Jégu & Santos 2000 | ***Serrasalmus altispinis 1*** | **ROM 97979** | **t18869** | | **Suriname** | **Maroni (Marowijne)** |
| 177 | *Serrasalmus altispinis* Merckx, Jégu & Santos 2000 | ***Serrasalmus altispinis 2*** | **ROM 97979** | **t18871** | | **Suriname** | **Maroni (Marowijne)** |
| 178 | *Serrasalmus altuvei* Ramírez 1965 | ***Serrasalmus altuvei 1*** | **G. Ortí** | **HLF-247** | | **Venezuela** | **Orinoco: Rio Cinaruco** |
| 179 | *Serrasalmus altuvei* Ramírez 1965 | ***Serrasalmus altuvei 2*** | **ROM** | **t02549** | | **Venezuela** | **Orinoco: Casiquiare** |
| 180 | *Serrasalmus brandtii* Lütken 1875 | ***Serrasalmus brandtii 1*** | **G. Ortí** | **GO-408** | | **Brazil** | **São Francisco** |
| 181 | *Serrasalmus* cf. *gibbus* Castelnau 1855 | ***Serrasalmus cf gibbus 1*** | tba ROM | **MK-18-015** | | **locality unknown** | **pet trade** |
| 182 | *Serrasalmus hastatus* Fink & Machado-Allison 2001 | ***Serrasalmus hastatus 1*** | **G. Ortí** | **GOLABS3** | | **locality unknown** | **locality unknown** |
| 183 | *Serrasalmus compressus* Jégu, Leão & Santos 1991 | ***Serrasalmus compressus 1*** | **UMMZ 250776** | **UMMZ T00907** | | **Guyana** | **Essequibo: Rewa River** |
| 184 | *Serrasalmus compressus* Jégu, Leão & Santos 1991 | ***Serrasalmus compressus 2*** | **UMMZ 250827** | **UMMZ T00970** | | **Guyana** | **Essequibo: Rewa River** |
| 162 | *Pristobrycon* cf. *eigenmanni* (Norman 1929) | ***Serrasalmus cf eigenmanni 1*** | **ANSP 188683** | **6928** | **t1627** | **Suriname** | **Maroni: Lawa** |
| 161 | *Serrasalmus maculatus* Kner 1858 | ***Serrasalmus cf eigenmanni 2*** | **ANSP 196873** | **t2220** | **t10388** | **Brazil** | **Amazonas: Xingu** |
| 186 | *Serrasalmus eigenmanni* Norman 1929 | ***Serrasalmus eigenmanni 1*** | **ROM 97245** | **t18297** | | **Guyana** | **Berbice** |
| 187 | *Serrasalmus eigenmanni* Norman 1929 | ***Serrasalmus eigenmanni 2*** | **ROM 100579** | **t20167** | | **Suriname** | **Corantin** |
| 185 | *Serrasalmus eigenmanni* Norman 1929 | ***Serrasalmus eigenmanni 3*** | **ROM 86351** | **t06269** | | **Guyana** | **Essequibo: Rupununi** |
| 188 | *Serrasalmus elongatus* Kner 1858 | ***Serrasalmus elongatus 1*** | **ANSP 196874** | **t2215** | **ANSP t10383** | **Brazil** | **Amazonas: Xingu** |
| 189 | *Serrasalmus elongatus* Kner 1858 | ***Serrasalmus elongatus 2*** | **ROM 98848** | **t19528** | | **Peru** | **Amazonas: Nanay** |

| | | | | | | | |
|---|---|---|---|---|---|---|---|
| 190 | *Serrasalmus elongatus* Kner 1858 | *Serrasalmus elongatus 3* | ROM 98848 | t19529 | | Peru | Amazonas: Nanay |
| 191 | *Serrasalmus geryi* Jégu & Santos 1988 | *Serrasalmus geryi 1* | ROM 105321 | t04783 | | Brazil | Amazonas: Tocantins |
| 193 | *Serrasalmus gouldingi* Fink & Machado-Allison 1992 | *Serrasalmus gouldingi 1* | AUM 43730 | V5270 | AUFT 3045 | Venezuela | Río Negro : Río Amazonas |
| 194 | *Serrasalmus irritans* Peters 1877 | *Serrasalmus irritans 1* | AUM 54065 | T09042 | | Venezuela | Orinoco: Apure |
| 195 | *Serrasalmus maculatus* Kner 1858 | *Serrasalmus maculatus 1* | ROM 103433 | t24153 | | Uruguay | Rio Uruguay |
| 196 | *Serrasalmus maculatus* Kner 1858 | *Serrasalmus maculatus 2* | ROM 103433 | t24155 | | Uruguay | Rio Uruguay |
| 199 | *Serrasalmus manueli* (Fernández-Yépez & Ramírez 1967) | *Serrasalmus manueli 1* | AUM 43731 | V5269 | AUFT 3054 | Venezuela | Casiquiare |
| 197 | *Serrasalmus manueli* (Fernández-Yépez & Ramírez 1967) | *Serrasalmus manueli 2* | ANSP 198074 | t3446 | t12381 | Brazil | Amazonas: Xingu |
| 198 | *Serrasalmus manueli* (Fernández-Yépez & Ramírez 1967) | *Serrasalmus manueli 3* | ANSP 198074 | t3457 | t12391 | Brazil | Amazonas: Xingu |
| 200 | *Serrasalmus medinai* Ramírez 1965 | *Serrasalmus medinai 1* | AUM 54056 | t09777 | | Venezuela | Orinoco: Ventuari |
| 201 | *Serrasalmus medinai* Ramírez 1965 | *Serrasalmus medinai 2* | ROM | t9710 | | Venezuela | Orinoco: Ventuari |
| 202 | *Serrasalmus rhombeus* (Linnaeus 1766) | *Serrasalmus rhombeus 1* | ANSP 195066 | t0606 | t8251 | Brazil | Amazonas: Xingu |
| 203 | *Serrasalmus rhombeus* (Linnaeus 1766) | *Serrasalmus rhombeus 2* | PUJ | | | Colomiba | Orinoco: Vaupes |
| 205 | *Serrasalmus rhombeus* (Linnaeus 1766) | *Serrasalmus rhombeus 3* | ROM | t20168 | | Suriname | Corantijn |
| 204 | *Serrasalmus rhombeus* (Linnaeus 1766) | *Serrasalmus rhombeus 4* | ROM 96128 | t16619 | | Guyana | Essequibo: Kuyuwini |
| 206 | *Serrasalmus serrulatus* (Valenciennes 1850) | *Serrasalmus serrulatus 1* | G. Ortí | GOLAB-224 | | Brazil | Rio Solimoes |
| 207 | *Serrasalmus spilopleura* Kner 1858 | *Serrasalmus spilopleura 1* | G. Ortí | SW2548 | | Brazil | Amazonas: Rio Negro |
| 208 | *Tometes ancylorhynchus* Andrade, Jégu & Giarrizzo 2016 | *Tometes ancylorhynchus 1* | tba ROM | | | Brazil | Amazonas: Jari |
| 209 | *Tometes ancylorhynchus* Andrade, Jégu & Giarrizzo 2016 | *Tometes ancylorhynchus 2* | tba ROM | | | Brazil | Amazonas: Jari |
| 210 | *Tometes kranponhah* Andrade, Jégu & Giarrizzo 2016 | *Tometes kranponhah 1* | ANSP 193019 | B2042 | t5841 | Brazil | Amazonas: Xingu: Iriri |
| 211 | *Tometes kranponhah* Andrade, Jégu & Giarrizzo 2016 | *Tometes kranponhah 2* | ROM | t15496 | | Brazil | Amazonas: Xingu: Iriri |
| 212 | *Tometes lebaili* Jégu, Keith & Belmont-Jégu 2002 | *Tometes lebaili 1* | ANSP 187099 | 6983 | t1633 | Suriname | Maroni: Lawa |
| 213 | *Tometes lebaili* Jégu, Keith & Belmont-Jégu 2002 | *Tometes lebaili 2* | ANSP 188681 | 7022 | t1634 | Suriname | Maroni: Litanie |
| 214 | *Tometes trilobatus* Valenciennes 1850 | *Tometes trilobatus 1* | UMMZ 250855 | UMMZ T00971 | | Guyana | Essequibo: Rewa River |
| 215 | *Utiaritichthys* cf. *longidorsalis* Jégu, de Morais & Santos 1992 | *Utiaritichthys aff longidorsalis 1* | ANSP 180811 | P4365 | t611 | Venezuela | Casiquiare |
